# Loss of pseudouridine synthases in the RluA family causes hypersensitive nociception in *Drosophila*

**DOI:** 10.1101/831917

**Authors:** Wan Song, W. Daniel Tracey

## Abstract

Nociceptive neurons of *Drosophila melanogaster* larvae are characterized by highly branched dendritic processes whose proper morphogenesis relies on a large number of RNA-binding proteins. Post-transcriptional regulation of RNA in these dendrites has been found to play an important role in their function. Here, we investigate the neuronal functions of two putative RNA modification genes, *RluA-1* and *RluA-2*, which are predicted to encode pseudouridine synthases. *RluA-1* is specifically expressed in larval sensory neurons while *RluA-2* expression is ubiquitous. Nociceptor-specific RNAi knockdown of *RluA-1* caused hypersensitive nociception phenotypes, which were recapitulated with genetic null alleles. These were rescued with genomic duplication and nociceptor-specific expression of *UAS-RluA-1-cDNA*. As with *RluA-1, RluA-2* loss of function mutants also displayed hyperalgesia. Interestingly, nociceptor neuron dendrites showed a hyperbranched morphology in the *RluA-1* mutants. The latter may be a cause or a consequence of heightened sensitivity in mutant nociception behaviors.

**Author Summary:** Pseudouridine (Psi) is a C5-glycoside isomer of uridine and it is the most common posttranscriptional modification of RNAs, including noncoding tRNAs, rRNAs, snRNAs as well as mRNAs. Although first discovered in the 1950s, the biological functions of Psi in multicellular organisms are not well understood. Interestingly, a marker for sensory neurons in *Drosophila* encodes for a putative pseudouridine synthase called RluA-1. Here, we report our characterization of nociception phenotypes for larvae with RluA-1 loss of function along with that of a related gene RluA-2. Disrupting either or both RluA-1 and RluA-2 resulted in hypersensitive nociception. In addition, RluA-1 mutants have more highly branched nociceptor neurites that innervate the epidermis. Our studies suggest an important role for the RluA family in nociception. This may be through its action on RNAs that regulate neuronal excitability and/or dendrite morphogenesis.

## Introduction

Pain serves an indispensable, protective role but when pain becomes pathological it can have a debilitating impact on human life. The total annual cost of pain to society in the United States was estimated by the Institute of Medicine to be up to $635 billion, which is greater than that of heart disease, cancer, and diabetes combined (Gaskin and Richard 2012). It is therefore of urgent importance to uncover the basic molecular and cellular mechanisms involved in pain in order to better treat it. Laboratory animal models of pain and nociception have played an essential role in identifying such mechanisms. *Drosophila* larvae respond to noxious thermal and mechanical stimuli through stereotyped rolling escape locomotion (in which the larva rotates around its long body axis), which is robust and easily distinguishable from other forms of locomotion (Tracey, Wilson et al. 2003). Rolling is used as a behavioral defense against attacks by parasitoid wasps which lay their eggs within the fly larvae using a sharp ovipositor (Hwang, Zhong et al. 2007). Successful wasp infection is lethal to the larvae and is a strong selective pressure in natural populations of *Drosophila.* Combined with the unparalleled genetic tools available for *Drosophila melanogaster*, this behavioral readout provides an excellent system to study the genetics of nociception and pain (Tracey, Wilson et al. 2003, Caldwell and Tracey 2010, Im and Galko 2012, Milinkeviciute, Gentile et al. 2012, Tracey 2017, Khuong, Wang et al. 2019). Previous studies have demonstrated a specific subset of dendritic arborization (da) sensory neurons in the peripheral nervous system, Class IV multidendritic da (cIVda) neurons, are of critical importance for thermal, mechanical and high intensity light nociception (Grueber, Jan et al. 2002, Hwang, Zhong et al. 2007, Xiang, Yuan et al. 2010). Further evidence suggests a lesser but significant contribution of Class II (cIIda) and Class III da (cIIIda) neurons in mechanical nociception (Hwang, Zhong et al. 2007, Kim, Coste et al. 2012, Hu, Petersen et al. 2017). In addition, great progress has been made in identifying the circuits in the larval abdominal ganglion that are involved in rolling escape locomotion (Ohyama, Jovanic et al. 2013, Ohyama, Schneider-Mizell et al. 2015, Chin and Tracey 2017, Burgos, Honjo et al. 2018).

Genes that are specifically expressed in multidendritic (md) neurons have been found to play a role in nociception. For instance, *painless* (Tracey, Wilson et al. 2003) is required for mechanical and thermal nociception and it is expressed in all four classes of md neurons. Similarly, *straightjacket* (Neely, Hess et al. 2010) is expressed in md neurons and required for avoidance of noxious heat. Mechanical nociception genes such as *pickpocket* (Zhong, Hwang et al. 2010), *ppk26*/*balboa* (Gorczyca, Younger et al. 2014, Mauthner, Hwang et al. 2014) and the polymodal nociception gene *dTRPA1-C/D* (Zhong, Bellemer et al. 2012) each show very specific expression in the cIVda neurons. Forward genetic screens have also identified a set of genes with enriched expression in the cIVda neurons which either inhibit or activate nociceptive pathways (Honjo, Mauthner et al. 2016).

The historically first known genetic marker with specific expression pattern in the da neurons was the *lacZ* enhancer trap *E7-2-36* (Brewster and Bodmer 1995). Later studies reported that this enhancer trap gene was inserted upstream of the *RluA-1* gene and DNA sequences from upstream of *RluA-1* caused expression of GAL4 in multidendritic neurons (Wang, Lo et al. 2011). The *RluA1* gene encodes an enzyme that is predicted to have pseudouridine synthase activity, but how this RNA modifying protein is involved in the function of nociceptive multidendritic neurons remains unknown.

A widespread importance of RNA-binding proteins in cIVda neuron dendrite morphogenesis and function was found in a recent large-scale RNAi screen that identified 88 genes encoding RNA-binding proteins whose knockdown caused aberrant dendrite morphogenesis (Olesnicky, Killian et al. 2014). The elaborate dendrite arbors of cIVda neurons project long distances from the neuronal cell body and mRNA granules are trafficked to these distant sites where they may undergo local translation. Indeed, RNA granules containing Nanos (Nos), Pumilio (Pum), Oskar (Osk), Fragile X Mental Retardation (FMRP) and other proteins have been shown to regulate the formation of higher order dendrites in these cells (Pan, Zhang et al. 2004, Ye, Petritsch et al. 2004, Brechbiel and Gavis 2008, Bianco, Dienstbier et al. 2010, Xu, Brechbiel et al. 2013). Interestingly, the *nos* mRNA itself is transported into dendrites via a pathway that relies on Oskar and Rumplestiltskin, two proteins that also play a role regulating *nos* in the early embryo (Xu, Brechbiel et al. 2013). Mechanical nociception defects are also observed in animals with disruption in these pathways (Xu, Brechbiel et al. 2013).

Pseudouridylation is the most common post-transcriptional RNA modification. Pseudouridine (Ψ), the C5-glycoside isomer of uridine, was initially found in many positions in rRNA, tRNA and snRNA in all organisms that have been investigated (Ge and Yu 2013). RNA-seq based global pseudouridine profiling has shown the presence of Ψ in many mRNAs and a large number of those sites were found to be dynamically regulated in yeast and human cells (Carlile, Rojas-Duran et al. 2014, Lovejoy, Riordan et al. 2014, Schwartz, Bernstein et al. 2014, Khoddami, Yerra et al. 2019). Dysfunctional pseudouridylation has been linked to several human diseases (Knight, Heiss et al. 1999, Fujiwara and Harigae 2013, de Brouwer, Abou Jamra et al. 2018). Since many sites of pseudouridylation in different organisms are evolutionarily conserved, *Drosophila melanogaster* may provide an excellent and genetically tractable metazoan system to elucidate some of these functions.

The isomerization of uridine to pseudouridine is catalyzed by six families of pseudouridine synthases. They function either as guide RNA directed ribonucleoprotein complexes or as stand-alone proteins (Koonin 1996, Hammal and Ferre-D’Amare 2006). In the *Drosophila* genome, 9 proteins have been identified with annotated pseudouridine synthase domains. Minifly (mfl), the RNA-dependent pseudouridine synthase homolog of human dyskerin (mouse NAP57 and yeast Cbf5), is required for somatic stem cell homeostasis and is essential for *Drosophila* viability and fertility (Phillips, Billin et al. 1998, Giordano, Peluso et al. 1999, Vicidomini, Petrizzo et al. 2017). Knockout of *Drosophila* Pus7, the human and yeast Pus7 homolog, results in increased aggressiveness in adult flies (de Brouwer, Abou Jamra et al. 2018). The function and specificity of other predicted pseudouridine synthases are largely unknown. Among the six families, the RluA family, which does not rely on guide RNAs, appears to be the most complex based on divergent substrate specificities in bacteria and yeast (Hoang, Chen et al. 2006). RluA family members in bacteria are involved in ribosomal assembly and growth (Raychaudhuri, Niu et al. 1999, Gutgsell, Deutscher et al. 2005) but their function in multicellular organisms has not been studied. Although pseudouridine synthases appear to function ubiquitously, as noted above, *Drosophila RluA-1*, a member in RluA family, has been reported to be specifically expressed in md neurons (Wang, Lo et al. 2011). Thus, we have investigated the role for *RluA-1* and its paralog *RluA-2*, in nociception pathways that are known to depend on md neurons. Our results indicate an important function for *RluA-1* and *RluA-2* in the regulation of nociception.

## Results

### *RluA-1* is expressed in the multidendritic neurons of the peripheral nervous system and in the cells of brain

Previously described reporter genes for *RluA-1* showed specific expression in larval multidendritic neurons (Wang, Lo et al. 2011). We replicated this finding by generating an *RluA-1^GAL4^* driver at the endogenous gene locus through recombination mediated cassette exchange (RMCE) of *RluA-1^MI06897^* (an intronic MiMIC) and a Trojan exon cassette (Diao, Ironfield et al. 2015). An mCD8GFP reporter driven by *RluA-1^GAL4^* was expressed in peripheral sensory neurons in each segment of the larval body wall (**Figure 1A**). In the dorsal cluster, GFP-positive signals were clearly detected in all four classes of md-da sensory neurons, dorsal multiple dendrite neuron (dmd1), external sensory (ES) and dorsal bipolar dendritic (dbd) neurons (**Figure 1B**). In the larval ventral nerve cord, the GFP-positive signals were seen in the axonal projections of the sensory neurons (**Figure 1C**). GFP signal was also observed in the unidentified clusters of neurons in the larval brain (**Figure 1C**). In the adult brain, significant signals were detected in the optic lobes and other small cell clusters of the central brain (**Figure S1A**).

**Figure 1.**
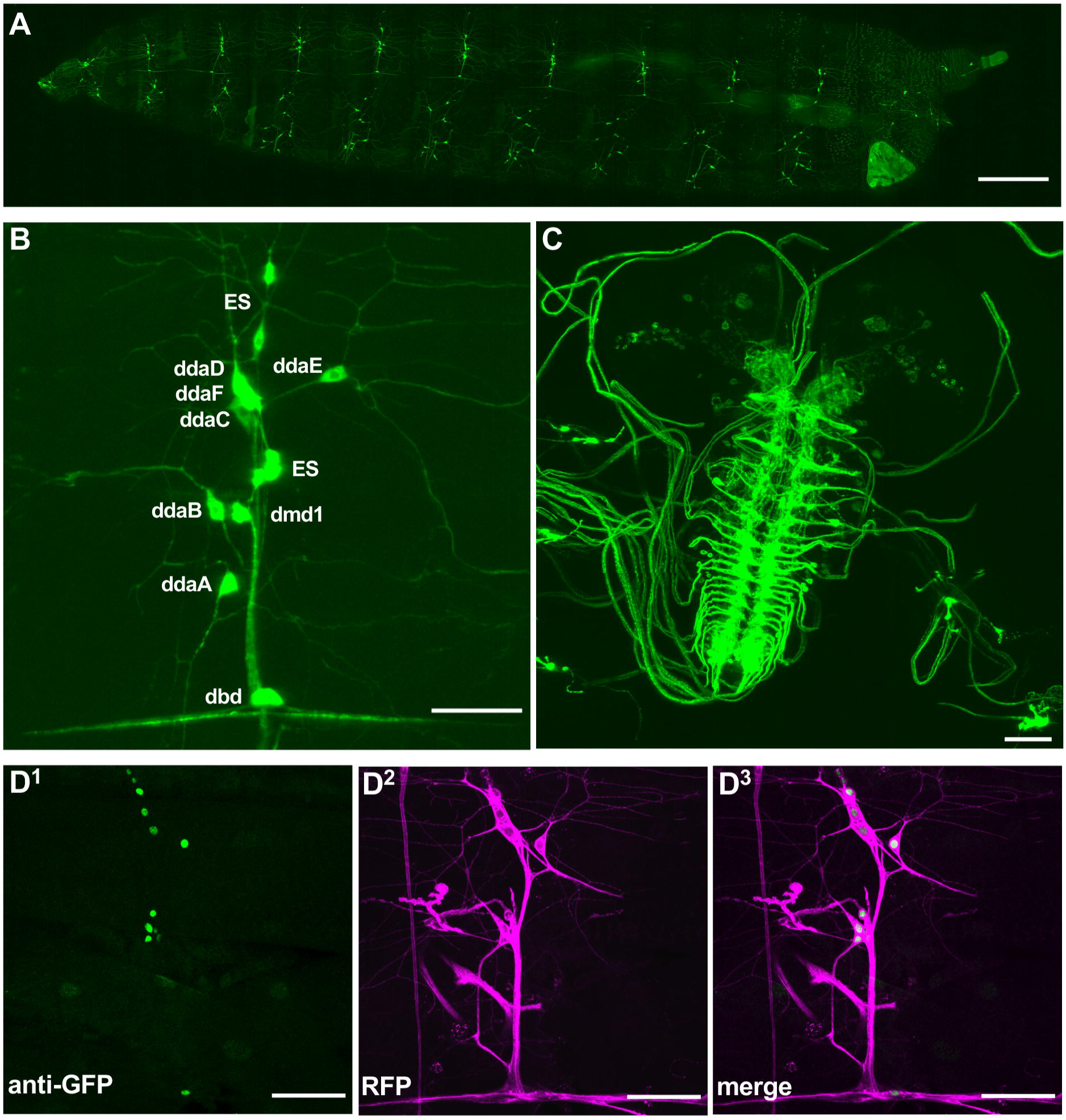
*RluA-1* gene and protein expression in *Drosophila melanogaster*. (**A**) A low magnification confocal micrograph of *RluA-1* gene expression pattern in the larval PNS (third instar *w^1118^; RluA-1^Gal4^*/*40xUAS-mCD8::GFP*). Note that GFP positive signals are detected in peripheral sensory neurons in each segment along the larval body wall. Scale bar=500μm. (**B**) Higher magnification of an abdominal dorsal PNS cluster. *RluA-1^Gal4^* is expressed in the cell body and dendrites of all classes of multidendritic neurons, external sensory (ES), dorsal multiple dendrite neuron (dmd1) and dorsal bipolar dendritic (dbd) neurons (ddaD and ddaE (Class I), ddaB (Class II), ddaA and ddaF (Class III), and ddaC (ClassIV), ES, dmd1 and dbd neurons are labeled). Scale bar = 50μm. (**C**). *RluA-1^Gal4^/+* driving expression of *40xUAS-mCD8::GFP/+* in the larval CNS. Labelling is observed in axonal projections of sensory neurons in the larval ventral nerve cord and unidentified clusters of neurons in the larval brain. Scale bar =5 0μm. (**D^1-3^**) anti-GFP immunohistochemistry of a third instar larval fillet preparation of *RluA-1-GFSTF/ Gal4109(2)80*>*UAS-mCD8-RFP*, immunoreactive signals are detected in the nuclei of multi-dendritic neurons (D^1^ green, GFP) surrounded by the membrane-localized RFP signal (D^2^ magenta, RFP) and merged image in D^3^. Scale bar = 50μm.

To determine the localization of the RluA-1 proteins, we generated a GFSTF exon trap (Nagarkar-Jaiswal, DeLuca et al. 2015) that expresses an in frame GFP fusion with *RluA-1* at the endogenous genomic locus (with RMCE of the *RluA-1^MI06897^* MiMIC element) (Nagarkar-Jaiswal, DeLuca et al. 2015). Although live imaging did not detect the EGFP tagged RluA-1 protein, immunostaining with anti-GFP labelled nuclei of larval multidendritic neurons, ES and dbd neurons also expressing a membrane-localized RFP (*Gal4109(2)80* > *UAS-CD8-RFP*, **Figure 1D**). The tagged RluA-1 protein was also detected in the cell bodies of neurons in optic lobes and other yet-to-be identified cells in the adult brain (**Figure S1B**).

### Reducing or removing *RluA-1* results in a hypersensitive thermal nociception phenotype

Given our confirmation of the expression of the *RluA1* gene and protein in the multidendritic neurons we tested for potential roles of RluA-1 in the regulation of nociception. To do so, we first performed tissue-specific knockdown using GAL4/UAS based RNA interference (RNAi). A cIVda specific driver (*ppk1.9-GAL4 UAS-dicer2*) (Ainsley, Pettus et al. 2003) was employed to drive an *RluA-1* RNAi construct in the larval cIVda nociceptors. We assessed potential insensitive phenotypes (with a probe temperature of 46°C) and potential hypersensitive phenotypes (with a temperature of 42°C) as previously described (Honjo, Mauthner et al. 2016). When stimulated with the higher temperature 46°C probe the *RluA-1-RNAi* larvae responded significantly faster than the *ppk-GAL4* driver alone controls. This genotype also responded faster than the UAS-RNAi controls but this difference was not statistically significant (**Figure 2A**). The results suggested that reducing *RluA-1* in classIV neurons may have made the larvae more sensitive to noxious heat. Indeed, a clearly hypersensitive phenotype for the *RluA1-* RNAi animals was seen when testing with 42°C probe (**Figure 2A**). The average latency to roll in the *RluA-1* knock-down animals was significantly faster than the driver alone animals or the *UAS-RNAi* control animals. These data combined suggest that reducing the activity of *RluA-1* in the noxious heat-responsive clVda cells caused thermal hyperalgesia.

**Figure 2.**
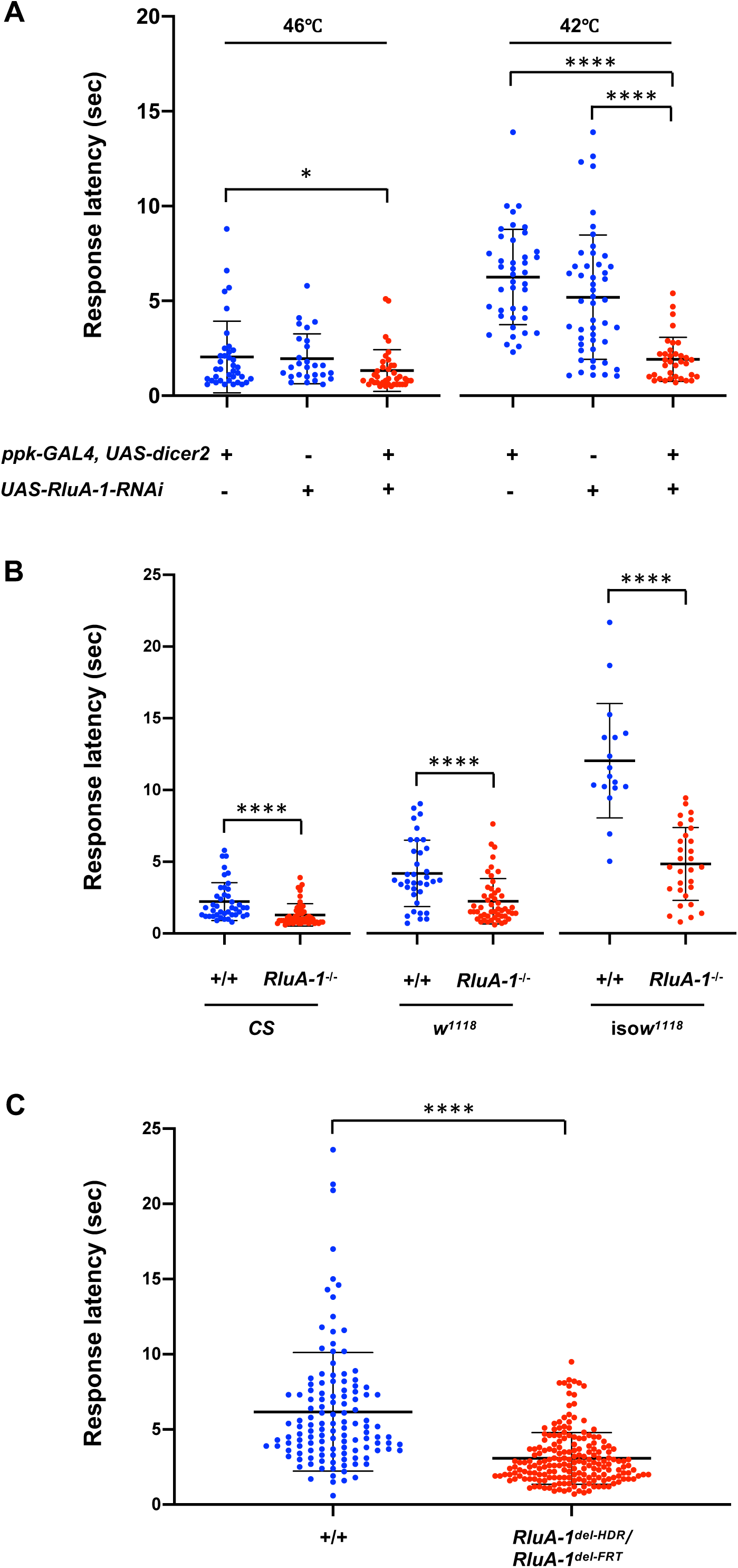
Thermal nociception in animals with *RluA-1* loss of function. **(A).** Class IV specific knock-down in *RluA-1* results in significant hypersensitive thermal nociception in larvae compared to the *ppk-GAL4* driver alone animals at 46℃ (average latency of 1.33 ± 1.10 sec for *ppk-Gal4* > *UAS-RluA-1-RNAi (*31719-R1), n=38 vs. 2.04 ± 1.89 sec for *ppk-GAL4* driver alone, n=37. P<0.05), albeit not statistically significant compared to the UAS-RNAi alone (1.96 ± 1.32 sec for UAS-RNAi alone, n=27). These differences are more pronounced at 42℃ (average latency of 1.92 ± 1.16 sec for *ppk-Gal4* > *UAS-RluA-1-RNAi*, n=35 vs. 6.25 ± 2.51 sec for *ppk-GAL4* driver alone controls, n=42 or 5.20 ± 3.28 sec for UAS-RNAi alone controls, n=49, P<0.0001). Kruskal-Wallis test with Dunn’s multiple comparison tests, only significant comparisons were labeled. **(B).** Homozygous null mutant *RluA-1^del-HDR^* larvae showed significantly faster response to noxious heat stimulation of 42°C compared to the corresponding control animals of *Canton-S* (average latency of 1.29 ± 0.78 sec in *RluA-1^-/-^*, n=54 vs. 2.22 ± 1.32 sec in *CS*, n=39), *w^1118^* (2.23 ± 1.57 sec in *RluA-1^-/-^*, n=51 vs. 4.16 ± 2.31 sec in *w^1118^*, n=36) and iso*w^1118^* (4.82 ± 2.53 sec in *RluA-1^-/-^*, n=30 vs. 11.99 ± 3.98 sec in iso*w^1118^*, n=17). Significance of comparisons are marked as **** (p<0.0001). Data were analyzed using Mann-Whitney non-parametric test. **(C).** Transheterozygote *RluA-1^del-HDR^*/*RluA-1^del-FRT^* showed hypersensitive thermal nociception responses compared to the controls (average latency of 3.07 ± 1.72 sec in *RluA-1^del-HDR^/RluA-1^del-FRT^*, n=195 vs. 6.17 ± 3.95 sec in +/+, n=125). The genetic background is *w^1118^* for *RluA-1^del-HDR^*, and iso*w^1118^* for *RluA-1^del-FRT^*. For the *RluA-1^del-HDR^/RluA-1^del-FRT^* transheterozygotes, data were pooled from the progeny from reciprocal crosses of *RluA-1^del-HDR^* to *RluA-1^del-FRT^*. To generate the control larvae, reciprocal crosses were made between the genetic background of *w^1118^* and iso*w^1118^* and the data from the progeny of these crosses were pooled. Significance of the comparison is marked as **** (exact p<0.0001). Data were analyzed using Mann-Whitney U test.

To further test the function of *RluA-1* we next generated a precise genetic deletion mutant of *RluA-1* in which 11.14 kbp including the entire *RluA-1* genomic region was removed by CRISPR-guided homologous recombination-directed repair (HDR) (Ran, Hsu et al. 2013). Homology arms of ∼1kb immediately flanking the CRIPSPR cleavage sites were used to direct the HDR (**Figure S2A**). The resultant deletion mutant (*RluA-1^del-HDR^*) was confirmed by PCR amplification and sequencing of PCR products from the targeted *RluA-1* locus (**Figure S2B and S2C**). To facilitate the behavioral comparisons, the *RluA-1^del-HDR^* mutant was backcrossed six times to our most commonly used genetic background strains Canton-S (CS), *w^1118^*, and isogenized *w^1118^* (iso*w^1118^*). As with class IV md neuron-specific RNAi knockdown, *RluA-1^del-HDR^* larvae showed significantly faster responses to noxious heat stimulation of 42°C compared to the corresponding control animals, regardless of genetic background (**Figure 2B**). Note that for unknown reasons, these genetic backgrounds vary in their baseline responses. This genetic background effect has a significant impact on the latency of larval nociception responses to noxious heat stimuli, with the most striking differences between homozygous *RluA-1^-/-^* and the relatively insensitive iso*w^1118^* background, followed by that of *w^1118^* background, and then the CS background (**Figure 2B**).

Given the sensitivity to genetic background, we performed an additional genetic test for the importance of *RluA-1*, by generating an independent mutant allele (*RluA-1^del-FRT^*) using FLP recombinase and FRT-bearing insertions (Parks, Cook et al. 2004) (*PBac{WH}^f02750^*(+) and *p{XP}^d2586^*(-), **Figure S2A**). Transheterozygous *RluA-1^del-FRT^*/*RluA-1^del-HDR^* mutant larvae displayed hypersensitivity to a 42°C stimulus (**Figure 2C**)) indicating that *RluA-1^del-FRT^* failed to complement *RluA-1^del-HDR^*. Failure of complementation of independently generated alleles created in distinct genetic backgrounds provides strong evidence that the hypersensitive nociceptive phenotypes observed are a consequence of the mutation of *RluA-1* and not due to unlinked mutations present in one background or another.

For the remainder of our behavioral studies, we relied on the isogenized *w^1118^* background. This had the advantage of greater genetic uniformity relative to *w^1118^* and CS, as well as showing the strongest hypersensitive mutant phenotype for *RluA-1*.

### Genetic rescue of *RluA-1* mutant restores thermal nociception response

To test that the mutation in *RluA-1* was the underlying cause of the hypersensitive nociception phenotype, we introduced a genomic rescue construct into the *RluA-1^del-HDR^* mutant background (a Bacterial Artificial Chromosome (BAC) (P6-D7)) (Venken, Carlson et al. 2009) covering the *RluA-1* gene region). With the 42°C stimulus, larvae with two copies of the duplication (Dp) in the background of *RluA-1^del-HDR^* showed rescue of the hypersensitivity (**Figure 3A**). This rescue with the genomic duplication was dosage dependent. Larvae with only one copy of the Dp in the *RluA-1^del-HDR^* background (*RluA-1^-/-^; Dp^+/-^*) responded more slowly than the mutant this difference was not statistically significant (**Figure 3A**). Combined, the results of genomic rescue experiments support the hypothesis that mutation of *RluA-1* is indeed the cause of the hypersensitive thermal nociception phenotype.

**Figure 3:**
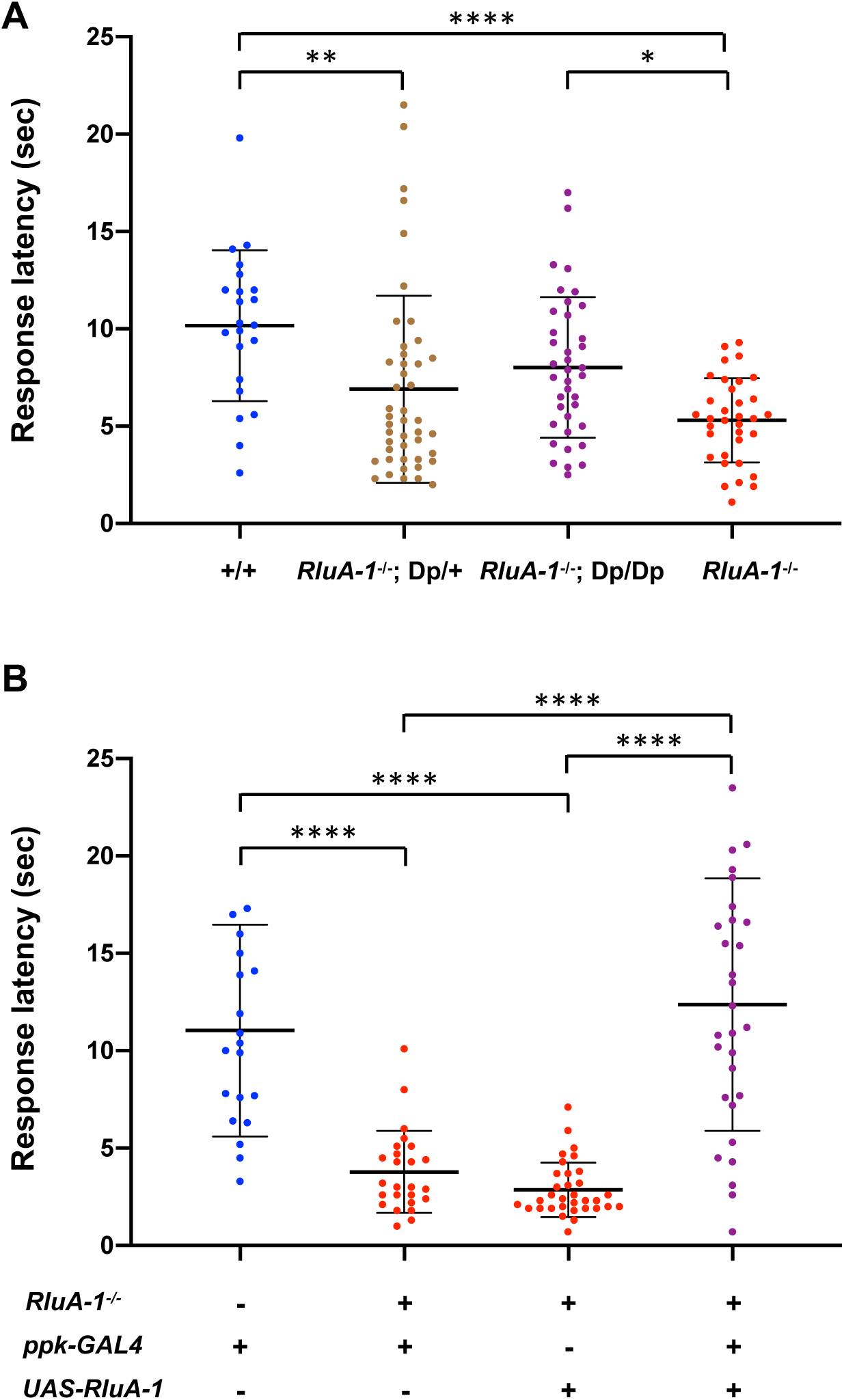
*RluA-1* nociception phenotypes with (A) chromosome duplication or (B) Class IV-specific expression of *UAS-RluA-1 cDNA*. (**A)**. The hypersensitive nociception phenotype with 42℃ thermal stimulus in *RluA-1^del-HDR^* null mutant was rescued by introducing a duplication (Dp) on the third chromosome which covers the *RluA-1* gene region (average latency of 8.02 ± 3.61 sec in *RluA-1^-/-^*; Dp/Dp, n=38, which is similar to the iso*w^1118^* genetic background (+/+, average latency of 10.16±3.87 sec, n=22), but significantly slower than the latency of 5.31 ± 2.16 sec in *RluA-1^-/-^*, n=34 (p=0.02)). Larvae with one copy of duplication showed slower response (average latency of 6.90 ± 4.81 sec in *RluA-1^-/-^; Dp/+*, n=45) without statistical significance compared to *RluA-1^del-HDR^*. Data were analyzed using Kruskal-Wallis test with Dunn’s multiple comparison tests. (**B).** The hypersensitive thermal nociception phenotype in *RluA-1^del-HDR^* larvae was completely reversed to that of the heterozygous *RluA-1* by Class IV specific expression of full length *RluA-1-cDNA* (*RluA-1^-/-^; ppk-Gal4^+/-^; UAS-RluA-1^+/-^*, average latency of 12.37 ±6.48 sec, n=30, is similar to *RluA-1^+/-^; ppk1.9-Gal4^+/-^*, average latency of 11.04 ± 5.44 sec, n=20 but significantly slower than either the driver alone (*RluA-1^-/-^; Ppk1.9-Gal4*^+/-^, average latency of 3.78±2.11 sec, n=25 or transgene alone controls (*RluA-1^-/-^; UAS-RluA-1*^+/-^, average latency of 2.86 ± 1.39 sec, n=33) at 42℃ thermal stimulus. Significance of the comparisons are marked as **** (p<0.0001). Data were analyzed using Kruskal-Wallis test with Dunn’s multiple comparison tests.

A caveat remained in that the genomic rescue construct included other genes in addition to *RluA-1.* Thus, genomic rescue did not rule out the possibility that a mutation tightly linked to *RluA-1,* but not in *RluA-1* itself, was responsible for the mutant phenotype. Thus, as a final test, we generated transgenic lines to express an *RluA-1* cDNA under the control of the GAL4/UAS system (*UAS-RluA-1*). Using the *UAS-RluA-1* we specifically restored *RluA-1* to cIVda neurons in the *RluA-1* null mutant background. When stimulated with the 42°C probe the animals with both the *ppkGal4* driver and the *UAS-RluA-1-cDNA* transgene in the *RluA-1 ^del-HDR^* showed a complete rescue from the hypersensitivity seen in the null mutant (**Figure 3B**). Neither the *ppkGal4* driver alone nor the *UAS-RluA-1* had an effect on the hypersensitive thermal nociception phenotype in the *RluA-1* null mutant background, excluding the possibility of non-specific effects of these transgenes (**Figure 3B**). In addition, overexpression of *UAS-RluA1* with a md neuron driver (MD-Gal4) had no effect on nociception behavior at 42°C stimulus ruling out the possibility that the increased latency seen in the rescue effect was a non-specific consequence of over-expression (**Figure S3**). Combined, these nociceptor-specific rescue experiments provide genetic confirmation that loss-of-function mutation in *RluA-1* causes hypersensitive thermal nociception and localizes the site of action for RluA-1 in this process to the nociceptors.

### *RluA-1* requirement for mechanosensory thresholds

The cIVda neurons are not only required for detection of noxious heat, they also contribute to sensing harsh mechanical stimulation (Hwang, Zhong et al. 2007, Zhong, Hwang et al. 2010, Mauthner, Hwang et al. 2014). Thus, we investigated the *RluA-1* mutant responses to noxious mechanical stimuli. When stimulated with a 30mN/720kPa Von Frey fiber, significantly more *RluA-1* ^del-HDR^ null mutant larvae performed the typical nociceptive rolling behavior compared to the iso*w^1118^* controls (**Figure 4A**). Even when the probe strength/pressure was reduced to 15mN/360kPa, the majority of *RluA-1^-/-^* larvae still rolled while less than half of control larvae rolled (**Figure 4A**), indicating the defect in *RluA-1* also caused hypersensitive mechanical nociception. Loss of *RluA-1* did not have any impact on behavioral responses to gentle touch (**Figure 4B**) suggesting a more specific involvement in nociception than for sensory processing in general.

**Figure 4:**
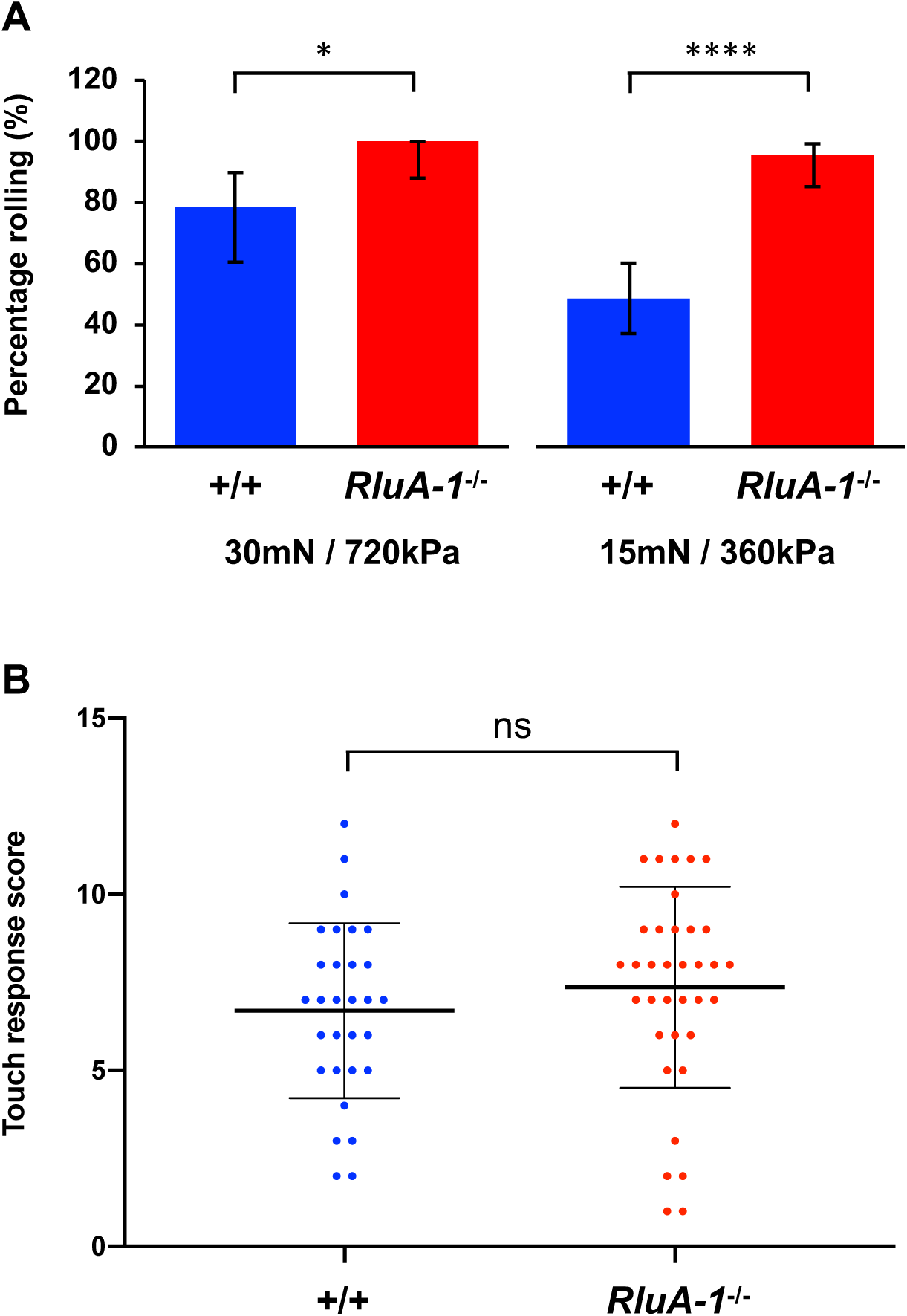
*RluA-1* mechanical nociception and gentle touch responses. (**A)**. In response to the noxious mechanical stimulus of 30mN/720kPa, all (100%) of the *RluA-1^del-HDR^* null mutant larvae (*RluA-1^-/-^*, n=28) rolled compared to the 78.57% of control larvae (iso*w^1118^* n=28) rolling. At the reduced stimulus of 15mN/360kPa, 95.56% of *RluA-1*^-/-^ (n=45) rolled while only 48.53% of control (n=68) rolled. Significance of the comparisons are marked as *(p<0.05) and **** (p<0.0001). Data were analyzed using Fisher’s exact test and presented as percentages ± 95% confidence intervals. (**B).** *RluA-1^del-HDR^* larvae (*RluA-1*^-/-^) had a gentle touch response score (6.70 ± 2.48, n=36) similar to the control larvae (iso*w^1118^,* 7.83 ± 2.69, n=30). ns, not significant (p>0.05). Data were analyzed with Student’s t test.

### Expression pattern and nociception functions for RluA-2

The *RluA-2* locus is adjacent to *RluA-1* on the second chromosome of *Drosophila melanogaster*. *RluA-2* has significant sequence similarity with *RluA-1* within the evolutionarily conserved pseudouridine synthase domain. This amino acid similarity of *RluA1* and *RluA2* suggested possible functional overlap for the encoded proteins. To investigate the expression of RluA-2 we generated a GFSTF line with RMCE of the *RluA-2^MI12981^* mimic element (**Figure S4A**) (Nagarkar-Jaiswal, DeLuca et al. 2015). Immunostaining with anti-GFP labelled nuclei in all of the cell types that we observed in third instar larvae (**Figure 5A**), including md neurons (**Figure 5B**).

**Figure 5:**
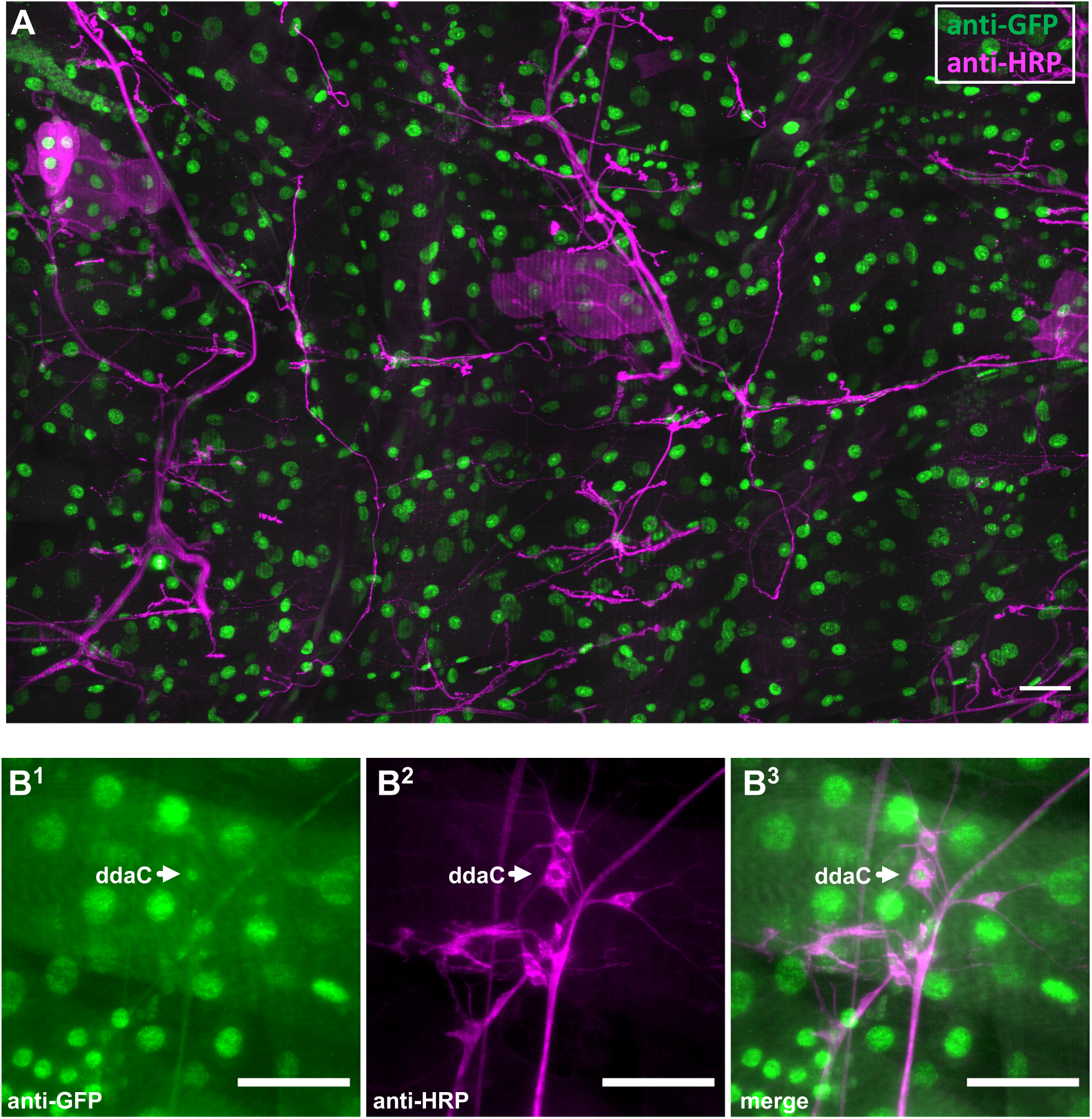
RluA-2 expression in *Drosophila* larvae. (**A).** In third instar larvae of *RluA-2-GFSTF* line, anti-GFP (green) immunoreactive signals are detected in the nuclei of all types of cells (neurons are labelled with anti-HRP (red)). Scale bar = 100μm. **(B).** In the larval abdominal dorsal PNS cluster of *RluA-2-GFSTF* line, GFP signal (green) is also detected in the nuclei of the md neurons, whose membranes are marked with anti-HRP (magenta). Arrow points to Class IV ddaC neuron. Scale bar = 50μm.

To test the function of *RluA-2* we generated a deletion (*RluA-2^del-HDR^*) to remove its pseudouridine synthase domain via CRISPR/Cas9 HDR (**Figure S4A**). Also, since RluA-1 and RluA-2 are both expressed in the md neurons (**Figure 1D** and **Figure 5B**), we generated a double mutant (*RluA-1^del-HDR^RluA-2^del-HDR^*), by injecting the *RluA-2^del-HDR^* constructs in the *RluA-1^del-HDR^* null mutant background (**Figure S4**). The double mutant allowed us to completely remove RluA pseudouridine synthase gene activity from the flies. Prior to functional assessment, the single mutant and the double mutant were backcrossed six times to the genetic background of iso*w^1118^*.

We tested each single mutant (*RluA-1^del-HDR^* and *RluA-2^del-HDR^*) together with the double mutant *RluA-1^del-HDR^ RluA-2^del-HDR^* side by side in thermal nociception assays with the 42°C thermal stimulus. *RluA-1^del-HDR^* larvae (*RluA-1^-/-^*) again responded significantly faster than the genetic background control (**Figure 6**). The *RluA-2^del-HDR^* single mutant larvae also displayed a faster response to the stimulus (**Figure 6**) and similar hypersensitivity was also seen in an independent allele for *RluA-2* that we generated by FLP/FRT mediated recombination (*RluA-2^del-FRT^*) allele (**Figure S5**). Finally, the double mutant *RluA-1^del-HDR^ RluA-2^del-HDR^* larvae showed a faster response than the control larvae (**Figure 6**) to the same extent as the single mutant of *RluA-2^del-HDR^*. These results indicated that *RluA-2*, like *RluA-1*, negatively regulates nociception. The finding that the double mutant did not show a more severe phenotype than either single mutant suggests that RluA-2 and RluA-1 have non-redundant functional roles, and that they may function in the same molecular pathway. When this pathway is disrupted, hypersensitive nociception results.

**Figure 6.**
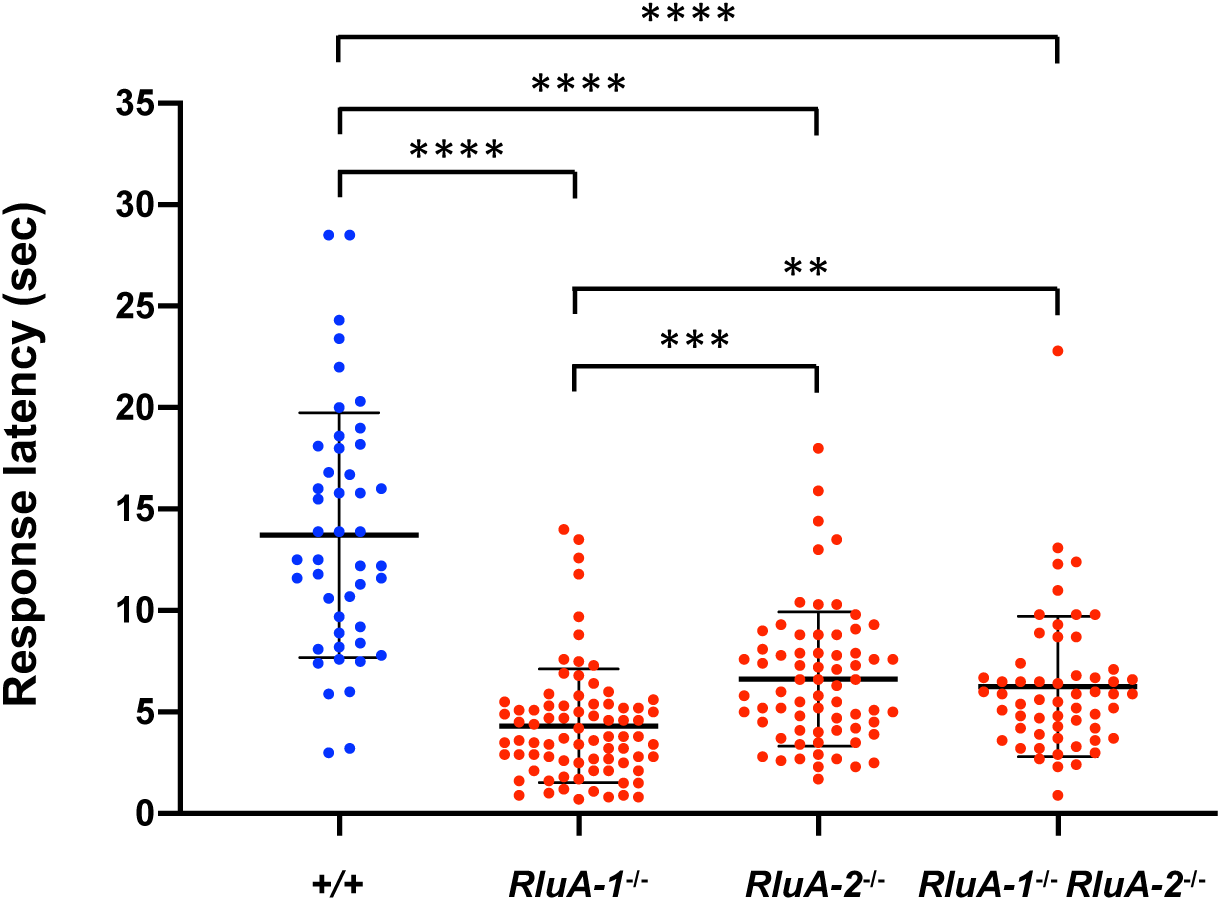
Thermal nociception responses of *RluA-1* and *RluA-2* single mutant, and *RluA-1 RluA-2* double mutant. Larvae of single mutant *RluA-1^del-HDR^* (*RluA-1^-/-^*), *RluA-2^del-HDR^* (*RluA-2^-/-^*), double mutant *RluA-1^del-HDR^ RluA-2^del-HDR^* (*RluA-1^-/-^ RluA-2^-/-^*) and the genetic background iso*w^1118^* (+/+) were stimulated with noxious heat probe of 42℃. *RluA-1^-/-^*, *RluA-2^-/-^*, and *RluA-1^-/-^ RluA-2^-/-^* all displayed faster responses to the stimulus compared to controls (*RluA-1^-/-^* average latency of 4.32 ± 2.80 sec, n= 79; *RluA-2^-/-^,* 6.63 ± 3.31 sec, n=68; (*RluA-1^-/-^RluA-2^-/-^*), 6.26 ± 3.46 sec, n=57; +/+, 13.72 ± 6.03 sec, n=46). Significance of comparisons are marked as ** (p<0.01), *** (p<0.001) or **** (p<0.0001). Data were analyzed using Kruskal-Wallis test with Dunn’s multiple comparison tests.

### *RluA-1* regulates neuronal dendrite morphology of nociceptors

A nociceptor-specific RNAi screen with thermal nociception assay discovered dozens of genes whose reduction caused either insensitive or hypersensitive thermal nociception (Honjo, Mauthner et al. 2016). Interestingly, some of those genes targeted with RNAi showed a reduced or increased branching of Class IV neuron dendrites. Reduced dendrite branching was often seen with nociceptive insensitivity while increased branching was found in some hypersensitive genotypes. Thus, regulation of Class IV neuron dendrite morphology is a commonly affected developmental pathway that is related to nociception phenotypes. Given this, we investigated the dendrite morphology of the cIVda neuron dendrites in the *RluA-1^del-HDR^* mutant. In mutant ddaC neurons visualized with *ppk-CD4-tdTom*ato, we observed a modest but significant increase in the number of dendrite branches (normalized by neuron size) and shorter average branch length in comparison to control animals (**Figures 7A and 7B**). We also found that dendritic branches in the *RluA-1^del-HDR^* ddaC neurons had higher frequency of isoneuronal cross-over events compared to the control (**Figures 7C and 7D**). This latter phenotype is suggestive of an isoneuronal tiling defect. Increased isoneuronal crossovers are also seen in mutants that affect dendrite attachment to the basal lamina (Han, Wang et al. 2012, Kim, Coste et al. 2012, Meltzer, Yadav et al. 2016, Tenenbaum, Misra et al. 2017). Whether or not these dendrite abnormalities play a causal role in the hypersensitive nociception phenotypes of the *RluA-1* mutant will be an interesting subject for future investigation.

**Figure 7.**
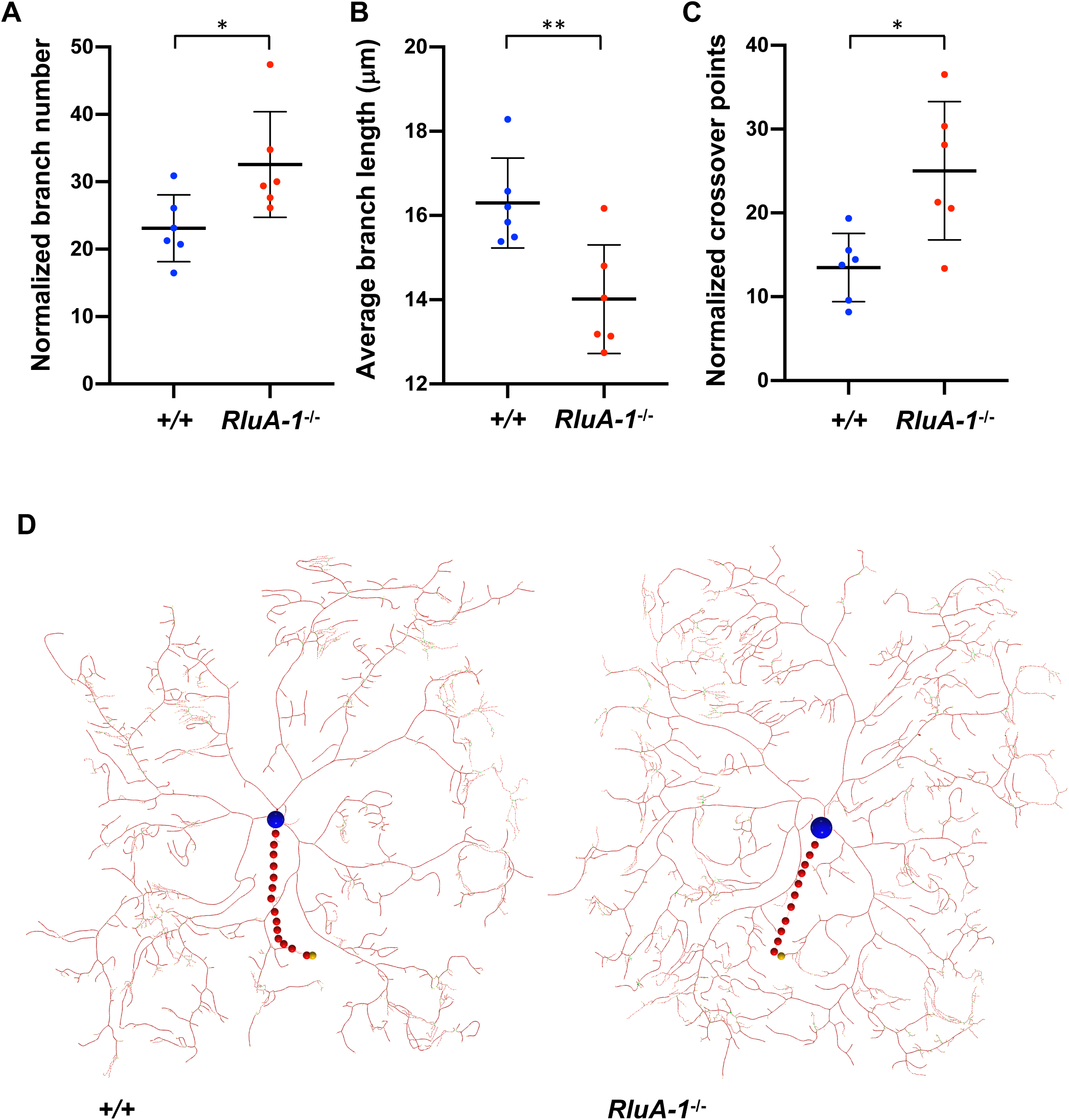
*RluA-1* ddaC neuron dendrite morphology. (**A-C).** Quantification of the branch number (**A**), average branch length (**B**), and isoneuronal cross-over points (**C**). (**A)**. Homozygous *RluA-1^del-HDR^* larvae showed a higher normalized branch number (total number of branches / (neuron size x 10^-4^μm^2^), 32.56 ± 7.83 in *RluA-1^-/-^*; *ppk-tdTom*, n=6 vs 23.10 ± 4.96 in *ppk-tdTom* control, n=6, p<0.05). (**B)**. The mean ddaC dendritic branch length in homozygous *RluA-1^del-HDR^* was shorter than that of the control (average branch length is 14.01 ± 1.29 μm in *RluA-1^-/-^; ppk-tdTom*, n=6 vs 16.30 ± 1.07 μm in *ppk-tdTom* control, n=6, p<0.01). (**C).** The number of ddaC dendritic isoneuronal crossovers in homozygous *RluA-1^del-HDR^* was higher compared to that of the control (normalized crossover points = crossover points / (neuron size x 10^-7^ μm^2^), 25.03 ± 8.24 in *RluA-1^-/-^*; *ppk-tdTom*, n=6 vs 13.48 ± 4.08 in *ppk-tdTom* control, n=6, p<0.05). Significant differences are marked as * (p<0.05) and ** (p<0.01). Data were analyzed with Student’s t test. (**D).** Representative traced dendritic structure of a class IV ddaC neuron of an L3 larva expressing *ppk-GAL4* > *UAS-td-Tom* in homozygous *RluA-1^del-HDR^* (*RluA-1^-/-^*, **left**) in comparison with that of the *ppk-tdTom* control (+/+, **right**). The position of the cell body is marked with blue circle and axon with red circles.

## Materials and methods

### Fly strains and husbandry

The following fly strains were obtained from Bloomington Stock Center: (*w^1118^*; *PBac{vas-Cas9}VK00027*), *Mi{MIC}RluA-1^[MI06897]^*, *Mi{MIC}RluA-2^[MI12981]^*, *Df(2L)Exel7048/CyO*, (*yw; Sp/CyO; pC-(lox-attB2-SA-T2A-Gal4-Hsp70)3*), (*yw hs-cre, vasΦC31; Sp/CyO; Sb/TM3Ser*), (*P{ry=hsFLP}, yw M{vas-int.B}ZH-2A; Sp/CyO*; *P{FRT-attB-{GFSTF}-attB(w+)-FRT}*), *ppk-CD4-tdTom*, *md-Gal4, UAS-mCD8::RFP*, *40XUAS-mCD8::GFP*. The following fly strains were obtained from Exelixis collection at Harvard Medical School: *PBac{WH}^f02750^*, *P{XP}d2586*, *PBac{WH}Grip75*^[f05483]^, *PBac{WH}RluA-2^[f07702]^*. The RNAi lines targeting *RluA-1* (31719-R1) were obtained from Kyoto Stock Center. The genetic duplication line (BAC ID: CH321-49P21) covering *RluA-1* (starting at 24,819,420 and ending at 24,910,132 on 2R) was obtained from Genetivision. The nos-Cas9 line used for generating *RluA-2^del-HDR^* and double mutant *RluA-1^del-HDR^RluA-2^del-HDR^* [*y sc v; {nos-Cas9} attP2* (TH00787.N)] was kindly provided by the Norbert Perrimon lab. All larvae used in experiments were reared on the Bloomington *Drosophila* medium in an incubator with controlled temperature (25°C) and humidity (70%) on a 12h light/12h dark cycle. Strains were otherwise maintained at room temperature.

### CRIPSR targeting of *RluA-1* and *RluA-2*

HDR for mutagenesis used the CRISPR/Cas9 (Gratz et al. 2013) to generate *RluA-1^del-HDR^*, *RluA-2^del-HDR^* and the double mutant *RluA-1^del-HDR^RluA-2^del-HDR^*. Target sites were selected using the flyCRISPR Optimal Target Finder. For the precise deletion of *RluA-1*, the gRNA target sites are 5’ end 5’-CCACTG|TGCAGCGGAAAATTCAC-3’ (where “|” represents the Cas9 cut site, which is 71bp upstream of the *RluA-1* transcriptional start) and 3’ end: 5’-ACATATATTCAAAAGCT|CTTTGG-3’ (cut site at 60bp downstream of *RluA-1* 3’UTR). We used the following primer pairs to amplify the 885bp *RluA-1* homology ARM1: *RluA-1*-ARM1F: 5’-AATACACCTGCATTATCGCTGGTCCCTGTGGCTTTGCAC-3’, *RluA-1*-ARM1R: 5’-AATACACCTGCAATTCTACCAGTGGGGCAAACCGCATTT-3’. We used *RluA-1*-ARM2F: 5’-GACTGCTCTTCGTATCTTTGGATGGTAAGTGCTTAAAC-3’ and *RluA-1*-ARM2R: 5’-ATTAGCTCTTCTGACCTTACAACTCCTTCAAGTCA-3’ to amplify 969 bp of *RluA-1* homology ARM2. For *RluA-2*, we used a target site at the 5’ end: 5’-GCAATCTATAGGTCTGC|GGAAGC-3’ (cut site at bp 5 of exon 3) and at the 3’ end: 5’AAGATAAACTACAGAGA|CCCCGG-3’ (cutting site in the middle of exon 8) so that the conserved pseudouridine synthase domain (located in exon 5) would be deleted. Primer pairs *RluA-2*-ARM1F: 5’CTAACACCTGCATATTCGCTCGAAACCCATTGTTAGCTG3’ and *RluA-2* ARM1R: 5’ CATTCACCTGCATTACTACGCAGACCTATAGATTGCAAT3’ were used to amplify the 979bp *RluA-2* homology ARM1. *RluA-2*-ARM2F: 5’CAGTGCTCTTCGTATCCCCGGCACAAAGGATCTCA-3’, and *RluA-2*-ARM2R: 5’ GACTGCTCTTCCGACATTTCAATGCCCTTGGCCAA-3’ were used to amplify the RluA-2 ARM2 (998bp). Rapid dsDNA donor cloning was carried out with the *pHD-DsRed-attP* vector (Beumer and Carroll 2014) and the guide RNA (gRNAs) were cloned into *pU6-BsbI-chiRNA* vector (Gratz, Cummings et al. 2013). Embryos of *vas-Cas9* on chromosome III (*w^1118^*; *PBac{y[+mDint2]=vas-Cas9}VK00027,* “injection line 1”) were injected with *RluA-1* dsDNA donor and gRNAs to generate G_0_ founders for *RluA-1^del-HDR^*. Embryos with *nos-Cas9* on chromosome III [*y sc v; {nos-Cas9} attP2* (TH00787.N), “injection line 2”] were injected with *RluA-2* dsDNA donor and gRNAs to create the G_0_ founders for *RluA-2^del-HDR^*. For generating the double mutant *RluA-1^del-HDR^RluA-2^del-HDR^*, *nos-Cas9* was first introduced to the *RluA-1^del-HDR^* mutant background and the DsRed marker in *RluA-1^del-HDR^* removed with CRE recombinase and the resultant homozygous strain [*yw*; *RluA-1^del-HDR^ΔDsRed*; *{nos-Cas9}attP2* (TH00787.N), “injection line 3”] was injected with the *RluA-2* dsDNA donor and gRNAs. All embryo injections of the dsDNA donor (at a concentration of 500ng/ul) and gRNAs (100ng/ul for each) were performed by the Model system Injections.

To identify the desired HDR mutants, G_0_ flies were crossed to *w^1118^* and single F_1_ founders were identified with DsRed fluorescence in the eyes (from *3XP3-DsRed* reporter) and mated with a second chromosome balancer strain to establish independent lines. Molecular testing for the desired events was performed by PCR on gDNA extracted from candidates placed over a deficiency (*Df(2L)Exelixis7048*) covering the *RluA-1* and *RluA-2* region (primer pairs marked in **Figures S2A and S4A, PCR amplication in Figure S2B and S4B**). PCR products sequenced across the *attp-loxP-3XP3-DsRed-SV40-loxP* fragment (**Figures S2C and Figures S4C**) confirmed accurate targeting of the loci. For behavioral analysis, the original deletion mutants were backcrossed to *CS*, *w^1118^* or iso*w^1118^* for six times. For each generation, five heterozygous females were selected for six successive backcrosses in vials and about 10 heterozygous females were used to cross to a second chromosome balancer to produce balanced mutant males in bottles and finally heterozygous mutant virgins and balanced males were crossed *en masse* in bottles to establish the balanced and homozygous mutant lines. In all crosses the DsRed fluorescence marker was used to follow the presence of the mutant.

### Generation of deletion line in *RluA-1* and *RluA-2* using FRT-mediated deficiency

FRT bearing insertions *{WH+}^f02750^* and *{XP-}^d2586^* and in *RluA-1* (locations marked in **Figure S2A**) were used for generating a deficiency allele *RluA-1^del-FRT^*. Insertions of *{WH-}RluA-2^F07702^* and *{WH-}Grip75^f05483^* (locations marked in **Figure S4A**) were used for generating the deficiency allele *RluA-2^del-FRT^*. Crossing and heat-shock schemes followed (Parks, Cook et al. 2004). Hybrid PCR with corresponding primers (WH5’ plus / XP5’ minus left and right primers) was used to screen for candidate lines with *w-* deletion in *RluA-1^del-FRT^* and two-sided PCR with left and right primers for WH3’ minus/ WH5’ minus was used for screening for candidate lines with *w+* deletion in *RluA-2^del-FRT^ (Parks, Cook et al. 2004).* Molecular testing for the deletion was performed by PCR on gDNA extracted from positive candidates placed over a deficiency (*Df(2L)Exelixis7048*) covering the *RluA-1* and *RluA-2* region. The primers used for PCR and sequencing verification in *RluA-1^del-FRT^* are *RluA-1-d2586*-up-F: 5’-AAAAATGCGGTTTGCCCC-3’, located upstream of the *{XP-}d2586* insertion site and *RluA-1-f02750*-down-R: 5’-AAGGGGTAACAAAAGTCCAG-3’, downstream of *{WH+}f02750 insertion site*.

### Generation of *RluA-1^GAL4^* using “Trojan-exon”

A triplet Trojan exon donor line on the 3rd chromosome *(yw; Sp/CyO; pC-(lox-attB2-SA-T2A-Gal4-Hsp70)3)* and an *RluA-1* insertion line containing an intronic MiMIC element (*Mi{MIC}RluA-1^[MI06897]^*) were used to generate the *RluA-1^Gal4^* driver using a crossing scheme as described by (Diao, Ironfield et al. 2015). Candidate males who have lost the *y^+^* selection marker associated with *MiMIC* were crossed to *40XUAS-mCD8::GFP* line and animals expressing GFP were selected to establish a stable line. The line with the correct linker (phase 0) was confirmed with sequencing of PCR products amplifying the left side with primers *RluA-1-*5494F: 5’-TGATGTTGCCCCATAACG-3’ and *T2A-Gal4*-Seq-1R: 5’ CGCTATCGATGCTCACGGTC-3’ and the right side with the primers *T2A-Gal4*-4F: 5’-ACACCGTGCTGATGCTGC-3’ and *RluA-1*-5907R: 5’-GAAAACATCGCACATCTGG-3’ of the *RluA-1* genome-*T2A-Gal4* insertion bordering region.

### Generation of GFSTF insertions in *RluA-1* and *RluA-2* by recombination mediated cassette exchange (RMCE)

Crossing, heat shocking and screening for EGFP tagged MiMIC lines in *RluA-1* and *RluA-2* were essentially carried out as described (Nagarkar-Jaiswal, DeLuca et al. 2015). Males carrying a MiMIC insertion in a coding intron of *RluA-1* (*RluA-1^MI06897^*) or *RluA-2* (*RluA-2^MI12981^*) were crossed to females carrying the *hs-FLP* and *vasa-phiC31* integrase on the X chromosome and a frame-specific (“phase 0”) *FRT* flanked multiple tag (*GFSTF*) cassette on chromosome III. Candidate males with mosaic *w-* eyes and *y*-bodies were individually crossed to *w; Sco/CyO; Sb/TM3 Ser* balancers to establish stocks. The presence and direction of the insertion were tested by PCR assays described (Venken, Schulze et al. 2011). Since the original MiMIC insertion in *RluA-1* or *RluA-2* and the respective gene are in the same orientation, positive PCR reaction 1 (with primers MiLF and TagR) and 4 (with primers MilR and TagF) as described (Venken, Schulze et al. 2011) indicated a successful RMCE event and resulted in expression.

### Generation of *UAS-RluA-1*

*Drosophila RluA-1* full length cDNA clone for *RluA-1* transcript A (FI04540) was obtained from the Drosophila Genome Resource Center (DGRC). The *UAS-RluA-1* expression constructs were generated with the ENTR/gateway system following the instructions of the manufacturer (Invitrogen). The *RluA-1*-cDNA-F (5’-CACCATGCAGAATTCTCCGGCT-3’, and *RluA-1*-cDNA+STOP-R (5’-TCATGCCGAGTCTAAGTG-3’ primers were used to amplify the open reading frame and cloned into *pENTR-D-TOPO* vector. The sequence was verified and then cloned to the P-element based destination vector *pTW*. Model System Injections performed injections into *w^1118^* embryos. F_0_ flies were crossed to *w^1118^* and single F_1_ founders were identified based on the *w*^+^ marker. Individual lines were mapped and balanced to establish stable stocks. A *UAS-RluA-1* line with relatively weak expression evaluated by Q-PCR in *RluA-1*-cDNA lines driven by *md-GAL4* was used for the cIV-neuron specific *RluA-1*-cDNA rescue experiment (**Figure 3B**).

### Larval behavioral analyses

Wandering 3rd instar larvae were washed out from vials and acclimated for 5 min in petri dishes before testing. Larval thermal nociception assays were conducted essentially as previously described (Caldwell and Tracey 2010, Mauthner, Hwang et al. 2014, Walcott, Mauthner et al. 2018), except that the probe is gently held against the lateral surface of abdominal segments 4, 5, or 6 until the animal completes a 360° roll along the dorsal-ventral axis. All animals tested eventually performed rolling and the response latency from all the animals was graphed for a given genotype. Larval mechanical nociception response assays were conducted as previously described (Mauthner, Hwang et al. 2014). Behavioral recording and scoring were performed with the observer blinded to the genotype.

For gentle touch assays, early L3 larvae were scooped out from the top layer of the fly food in the vials and 5-10 larvae were briefly rinsed with PBS and allowed to acclimate on 1% agarose in a plate for 5 min before testing with an eyelash fixed to the end of a paintbrush. Each larva was brushed with the eyelash on segments T1-A3 for 4 times and the responses were recorded and summarized using a gentle touch scale (Tsubouchi, Caldwell et al. 2012).

### Immunohistochemistry and microscopy

The following primary antibodies were used for immunofluorescence: rabbit anti-GFP (ab6556, Abcam, 1:500), mouse anti-GFP (ab38689, Abcam, 1:500), Rabbit anti-HRP (1:100), mouse anti-nc82 (DSHB, supernatant, 1:30). Alexa Fluor 488, 633 were used at 1:1000 as secondary antibodies. Detailed immunostaining protocol is available on request. Images for immunostained tissues were taken on a Zeiss LSM 5 LIVE confocal microscope using a 40X objective except for **Figure 1D** and **Figure S1B**, which were taken on a Zeiss LSM880 using the 63X objective.

### CIV dendrite imaging, tracing and analysis

To image class IV dendrites, the DsRed marker in *RluA-1^del-HDR^* was first removed with CRE recombinase (*w^1118^; RluA-1^del-HDRΔDsRed^*) and *ppk1.9-CD4::tdTom* was introduced to the mutant background and a stable homozygous stock was established. For CIV dendrite analysis, six virgins and three males were crossed in each vial for the mutant (*w^1118^*; *RluA-1^del-HDRΔDsRed^; ppk1.9-CD4::tdTom*) and control *(ppk1.9-CD4::-td-Tom*). Wandering larvae were anesthetized with diethyl ether in a sealed glass chamber for 15min before being arranged on a slide and covered with 50 mm glass coverslip. Neurons expressing the fluorescently tagged markers were visualized on a Zeiss LSM 5 Live confocal microscope with a 40X oil objective (Plan Apo M27, NA 0.8). Images were collected as 5×3 tiles scans of z-stacks with 512×512 resolution. A MatLab build was used for initial automatic tracing of the ddaC neuron dendrites from the confocal z-stack series TIFF images (Gulyanon 2016). The generated SWC files were overlaid onto the maximum intensity projected image of the neuron in neuTube (Feng, Zhao et al. 2015) and manually curated to eliminate tracing errors made by MatLab. The corrected images were then analyzed with MatLab to extract neuron features of interest including number of branches, average branch length, and neuron size (the estimated size of the neuron, defined as the area of the minimum bounding circle) (Gulyanon 2016). Iso-neuronal crossover events were quantified manually from the traced dendrites for each genotype (n=6 neurons).

### Statistical analysis

Statistics were performed using GraphPad Prism 4. Thermal and gentle touch behavioral data were compared with an unpaired non-parametric Mann-Whitney test when comparing two groups and Kruskal-Wallis test when comparing three or more groups. Mechanical nociception behavioral data were compared with Fisher’s exact test. Dendrite morphology data were compared with the Student’s t test. Error bars represent S.D. in all the figures unless otherwise specified.

### Data and Reagent Availability Statement

Strains and plasmids are available upon request. All supplementary figures have been uploaded to figshare. The authors affirm that all data necessary for confirming the conclusions of the article are present within the article and its figures.

## Discussion

Given the well-established nociceptive role of md-neurons, we have investigated the historically first known molecular marker for md-neurons in nociception pathways. This gene encodes the RluA-1 protein in the RluA family of pseudouridine synthases. Our studies clearly demonstrate that loss of function for either *RluA-1* or *RluA-2* produce hyperalgesia in third instar *Drosophila* larvae. Tissue-specific RNAi, genetic null mutant, and cDNA rescue experiments all indicate that loss of the *RluA-1* gene from whole animals, or specifically from nociceptors, results in hyperalgesia. A newly generated *RluA-1^GAL4^* driver showed specific expression in larval multidendritic and ES neurons. As well, a GFP exon trap for RluA-1 protein localized to the nuclei of these neurons. A small number of unidentified neurons in the larval brain were also revealed by *RluA-1^GAL4^* and we observed expression of *RluA-1^GAL4^* driven mCD8GFP and GFP tagged RluA-1 in cells of the adult brain, which included the optic lobe.

Although loss of *RluA-2* also caused hyperalgesia, its expression pattern was ubiquitous and included multidendritic neurons. Like the RluA-1 GFP exon trap, the RluA-2 GFP exon trap labelled nuclei. The nuclear localization for both RluA-1 and RluA-2 may indicate that these proteins act on RNA targets prior to export of the nucleus, or that they predominantly act upon nuclear localized RNAs. A caveat to this interpretation is that we have yet to demonstrate that these molecules function as genuine pseudouridine synthases, and this function remains a hypothesis that is based on the known function of evolutionary homologues. This hypothetical function requires future biochemical verification.

We also observed that *RluA-1* mutants showed an increase in the number of dendrite branches relative to control genotypes as well as an increase in isoneuronal crossovers. Transcription factors, cytoskeletal regulators, motor proteins, secretory pathways and cell adhesion molecules all function in concert to develop and maintain optimum dendrite morphology (Corty, Matthews et al. 2009). RluA-1 may modify RNAs for those genes that regulate dendritic morphology, or changes in the dendrite morphology could be an indirect consequence of neuronal sensitivity that is regulated by RluA-1. It is noteworthy that prior studies have noted a potential link between the degree of dendrite branching and the sensitivity of nociception behaviors (Honjo, Mauthner et al. 2016). In other cases, axonal factors such as the ion channel SK have been found to be important in regulating the cIVda neuron excitability (Onodera, Baba et al. 2017, Walcott, Mauthner et al. 2018). Whether the dendrite branching phenotype that we observe in *RluA-1* mutants is a cause or a consequence of hypersensitivity will be an interesting question for future studies.

A large body of literature indicates that RNA trafficking and local translation is important in dendrites and axons of neurons (Glock, Heumuller et al. 2017, Rangaraju, Tom Dieck et al. 2017). Relative to uridine the pseudourine base is believed to have enhanced rotational freedom which may alter conformation of RNA secondary structures. As well, an additional hydrogen bond donor present in pseudouridine may favor alternative base-pairing interactions in RNA. These properties may consequently alter RNA localization, stability and/or efficiency of translation (Arnez and Steitz 1994, Newby and Greenbaum 2002, Kariko, Muramatsu et al. 2008). Pseudouridines can also influence decoding during translation as pseudouridylation of nonsense codons has been shown to suppress translation termination both *in vitro* and *in vivo* (Karijolich and Yu 2011). Thus, another possible function for pseudouridylation in nociceptive neurons could be to favor read-through of pseudouridylated stop codons to generate novel sequences at protein carboxy termini.

The precise mechanism explaining the involvement of RluA proteins in nociception can only be elucidated by identifying the RNA targets of these enzymes. Several groups have developed methods using next generation sequencing methods to identify the pseudouridine sites in transcriptomes (Carlile, Rojas-Duran et al. 2014, Lovejoy, Riordan et al. 2014, Schwartz, Bernstein et al. 2014, Li, Zhu et al. 2015, Khoddami, Yerra et al. 2019). Future investigations applying these methods to wild type and *RluA* mutants in *Drosophila* will help us to identify the RluA targets and to further define the underlying mechanisms.

## Acknowledgements

The authors thank Drosophila Genomic Resource Center (supported by NIH grant 2P40OD010949) for clones. Stocks obtained from the Bloomington Drosophila Stock Center (NIH P40OD018537) were used in this study. We also thank the KYOTO Stock Center Kyoto in Kyoto Institute of Technology and Harvard Medical School for providing fly stocks used in this study. Members of Tracey lab provided helpful feedback on the manuscript and Jayce Brown Culbertson assisted with dendrite tracing.

## Notes

#### Summary of Updates

Presentation of figures revised

## References

Ainsley, J. A., J. M. Pettus, D. Bosenko, C. E. Gerstein, N. Zinkevich, M. G. Anderson, C. M. Adams, M. J. Welsh and W. A. Johnson (2003). “Enhanced locomotion caused by loss of the Drosophila DEG/ENaC protein Pickpocket1.” Curr Biol 13(17): 1557–1563.

Arnez, J. G. and T. A. Steitz (1994). “Crystal structure of unmodified tRNA(Gln) complexed with glutaminyl-tRNA synthetase and ATP suggests a possible role for pseudo-uridines in stabilization of RNA structure.” Biochemistry 33(24): 7560–7567.

Beumer, K. J. and D. Carroll (2014). “Targeted genome engineering techniques in Drosophila.” Methods 68(1): 29–37.

Bianco, A., M. Dienstbier, H. K. Salter, G. Gatto and S. L. Bullock (2010). “Bicaudal-D regulates fragile X mental retardation protein levels, motility, and function during neuronal morphogenesis.” Curr Biol 20(16): 1487–1492.

Brechbiel, J. L. and E. R. Gavis (2008). “Spatial regulation of nanos is required for its function in dendrite morphogenesis.” Curr Biol 18(10): 745–750.

Brewster, R. and R. Bodmer (1995). “Origin and specification of type II sensory neurons in Drosophila.” Development 121(9): 2923–2936.

Burgos, A., K. Honjo, T. Ohyama, C. S. Qian, G. J. Shin, D. M. Gohl, M. Silies, W. D. Tracey, M. Zlatic, A. Cardona and W. B. Grueber (2018). “Nociceptive interneurons control modular motor pathways to promote escape behavior in Drosophila.” Elife 7.

Caldwell, J. C. and W. D. Tracey (2010). “Alternatives to Mammalian Pain Models 2: Using Drosophila to Identify Novel Genes Involved in Nociception.” Analgesia: Methods and Protocols 617: 19–29.

Carlile, T. M., M. F. Rojas-Duran, B. Zinshteyn, H. Shin, K. M. Bartoli and W. V. Gilbert (2014). “Pseudouridine profiling reveals regulated mRNA pseudouridylation in yeast and human cells.” Nature 515(7525): 143-+.

Chin, M. R. and W. D. Tracey, Jr. (2017). “Nociceptive Circuits: Can’t Escape Detection.” Curr Biol 27(16): R796–R798.

Corty, M. M., B. J. Matthews and W. B. Grueber (2009). “Molecules and mechanisms of dendrite development in Drosophila.” Development 136(7): 1049–1061.

de Brouwer, A. P. M., R. Abou Jamra, N. Kortel, C. Soyris, D. L. Polla, M. Safra, A. Zisso, C. A. Powell, P. Rebelo-Guiomar, N. Dinges, V. Morin, M. Stock, M. Hussain, M. Shahzad, S. Riazuddin, Z. M. Ahmed, R. Pfundt, F. Schwarz, L. de Boer, A. Reis, D. Grozeva, F. L. Raymond, S. Riazuddin, D. A. Koolen, M. Minczuk, J. Y. Roignant, H. van Bokhoven and S. Schwartz (2018). “Variants in PUS7 Cause Intellectual Disability with Speech Delay, Microcephaly, Short Stature, and Aggressive Behavior.” Am J Hum Genet 103(6): 1045–1052.

Diao, F., H. Ironfield, H. Luan, F. Diao, W. C. Shropshire, J. Ewer, E. Marr, C. J. Potter, M. Landgraf and B. H. White (2015). “Plug-and-play genetic access to drosophila cell types using exchangeable exon cassettes.” Cell reports 10(8): 1410–1421.

Feng, L., T. Zhao and J. Kim (2015). “neuTube 1.0: A New Design for Efficient Neuron Reconstruction Software Based on the SWC Format.” eNeuro 2(1).

Fujiwara, T. and H. Harigae (2013). “Pathophysiology and genetic mutations in congenital sideroblastic anemia.” Pediatrics International 55(6): 675–679.

Gaskin, D. J. and P. Richard (2012). “The economic costs of pain in the United States.” J Pain 13(8): 715–724.

Ge, J. and Y. T. Yu (2013). “RNA pseudouridylation: new insights into an old modification.” Trends Biochem Sci 38(4): 210–218.

Giordano, E., I. Peluso, S. Senger and M. Furia (1999). “minifly, a Drosophila gene required for ribosome biogenesis.” J Cell Biol 144(6): 1123–1133.

Glock, C., M. Heumuller and E. M. Schuman (2017). “mRNA transport & local translation in neurons.” Curr Opin Neurobiol 45: 169–177.

Gorczyca, D. A., S. Younger, S. Meltzer, S. E. Kim, L. Cheng, W. Song, H. Y. Lee, L. Y. Jan and Y. N. Jan (2014). “Identification of Ppk26, a DEG/ENaC Channel Functioning with Ppk1 in a Mutually Dependent Manner to Guide Locomotion Behavior in Drosophila.” Cell Rep 9(4): 1446–1458.

Gratz, S. J., A. M. Cummings, J. N. Nguyen, D. C. Hamm, L. K. Donohue, M. M. Harrison, J. Wildonger and K. M. O’Connor-Giles (2013). “Genome engineering of Drosophila with the CRISPR RNA-guided Cas9 nuclease.” Genetics 194(4): 1029–1035.

Grueber, W. B., L. Y. Jan and Y. N. Jan (2002). “Tiling of the Drosophila epidermis by multidendritic sensory neurons.” Development 129(12): 2867–2878.

Gulyanon, S., Sharifai, N., Kim, M. D., Chiba, A., Tsechpenakis, G. (2016). CRF formulation of active contour population for efficient three-dimensional neurite tracing. 2016 IEEE 13th International Symposium on Biomedical Imaging: 593–597.

Gutgsell, N. S., M. P. Deutscher and J. Ofengand (2005). “The pseudouridine synthase RluD is required for normal ribosome assembly and function in Escherichia coli.” Rna-a Publication of the Rna Society 11(7): 1141–1152.

Hammal, T. and A. R. Ferre-D’Amare (2006). “Pseudouridine synthases.” Chemistry & Biology 13(11): 1125–1135.

Han, C., D. Wang, P. Soba, S. Zhu, X. Lin, L. Y. Jan and Y. N. Jan (2012). “Integrins regulate repulsion-mediated dendritic patterning of drosophila sensory neurons by restricting dendrites in a 2D space.” Neuron 73(1): 64–78.

Hoang, C., J. J. Chen, C. A. Vizthum, J. M. Kandel, C. S. Hamilton, E. G. Mueller and A. R. Ferre-D’Amare (2006). “Crystal structure of pseudouridine synthase RluA: Indirect sequence readout through protein-induced RNA structure.” Molecular Cell 24(4): 535–545.

Honjo, K., S. E. Mauthner, Y. Wang, J. H. Skene and W. D. Tracey, Jr. (2016). “Nociceptor-Enriched Genes Required for Normal Thermal Nociception.” Cell Rep 16(2): 295–303.

Hu, C., M. Petersen, N. Hoyer, B. Spitzweck, F. Tenedini, D. Wang, A. Gruschka, L. S. Burchardt, E. Szpotowicz, M. Schweizer, A. R. Guntur, C. H. Yang and P. Soba (2017). “Sensory integration and neuromodulatory feedback facilitate Drosophila mechanonociceptive behavior.” Nat Neurosci 20(8): 1085–1095.

Hwang, R. Y., L. Zhong, Y. Xu, T. Johnson, F. Zhang, K. Deisseroth and W. D. Tracey (2007). “Nociceptive neurons protect Drosophila larvae from parasitoid wasps.” Curr Biol 17(24): 2105–2116.

Im, S. H. and M. J. Galko (2012). “Pokes, sunburn, and hot sauce: Drosophila as an emerging model for the biology of nociception.” Developmental Dynamics 241(1): 16–26.

Karijolich, J. and Y. T. Yu (2011). “Converting nonsense codons into sense codons by targeted pseudouridylation.” Nature 474(7351): 395-+.

Kariko, K., H. Muramatsu, F. A. Welsh, J. Ludwig, H. Kato, S. Akira and D. Weissman (2008). “Incorporation of pseudouridine into mRNA yields superior nonimmunogenic vector with increased translational capacity and biological stability.” Mol Ther 16(11): 1833–1840.

Khoddami, V., A. Yerra, T. L. Mosbruger, A. M. Fleming, C. J. Burrows and B. R. Cairns (2019). “Transcriptome-wide profiling of multiple RNA modifications simultaneously at single-base resolution.” Proc Natl Acad Sci U S A 116(14): 6784–6789.

Khuong, T. M., Q. P. Wang, J. Manion, L. J. Oyston, M. T. Lau, H. Towler, Y. Q. Lin and G. Neely (2019). “Nerve injury drives a heightened state of vigilance and neuropathic sensitization in Drosophila.” Sci Adv 5(7): eaaw4099.

Kim, S. E., B. Coste, A. Chadha, B. Cook and A. Patapoutian (2012). “The role of Drosophila Piezo in mechanical nociception.” Nature 483(7388): 209–212.

Knight, S. W., N. S. Heiss, T. J. Vulliamy, S. Greschner, G. Stavrides, G. S. Pai, G. Lestringant, N. Varma, P. J. Mason, I. Dokal and A. Poustka (1999). “X-linked dyskeratosis congenita is predominantly caused by missense mutations in the DKC1 gene.” American Journal of Human Genetics 65(1): 50–58.

Koonin, E. V. (1996). “Pseudouridine synthases: four families of enzymes containing a putative uridine-binding motif also conserved in dUTPases and dCTP deaminases.” Nucleic Acids Res 24(12): 2411–2415.

Li, X. Y., P. Zhu, S. Q. Ma, J. H. Song, J. Y. Bai, F. F. Sun and C. Q. Yi (2015). “Chemical pulldown reveals dynamic pseudouridylation of the mammalian transcriptome.” Nature Chemical Biology 11(8): 592–U593.

Lovejoy, A. F., D. P. Riordan and P. O. Brown (2014). “Transcriptome-Wide Mapping of Pseudouridines: Pseudouridine Synthases Modify Specific mRNAs in S. cerevisiae.” Plos One 9(10).

Mauthner, S. E., R. Y. Hwang, A. H. Lewis, Q. Xiao, A. Tsubouchi, Y. Wang, K. Honjo, J. H. P. Skene, J. Grandl and W. D. Tracey (2014). “Balboa Binds to Pickpocket In Vivo and Is Required for Mechanical Nociception in Drosophila Larvae.” Current Biology 24(24).

Meltzer, S., S. Yadav, J. Lee, P. Soba, S. H. Younger, P. Jin, W. Zhang, J. Parrish, L. Y. Jan and Y. N. Jan (2016). “Epidermis-Derived Semaphorin Promotes Dendrite Self-Avoidance by Regulating Dendrite-Substrate Adhesion in Drosophila Sensory Neurons.” Neuron 89(4): 741–755.

Milinkeviciute, G., C. Gentile and G. G. Neely (2012). “Drosophila as a tool for studying the conserved genetics of pain.” Clinical Genetics 82(4): 359–366.

Nagarkar-Jaiswal, S., S. Z. DeLuca, P.-T. Lee, W.-W. Lin, H. Pan, Z. Zuo, J. Lv, A. C. Spradling and H. J. Bellen (2015). “A genetic toolkit for tagging intronic MiMIC containing genes.” eLife 4.

Neely, G. G., A. Hess, M. Costigan, A. C. Keene, S. Goulas, M. Langeslag, R. S. Griffin, I. Belfer, F. Dai, S. B. Smith, L. Diatchenko, V. Gupta, C. P. Xia, S. Amann, S. Kreitz, C. Heindl-Erdmann, S. Wolz, C. V. Ly, S. Arora, R. Sarangi, D. Dan, M. Novatchkova, M. Rosenzweig, D. G. Gibson, D. Truong, D. Schramek, T. Zoranovic, S. J. Cronin, B. Angjeli, K. Brune, G. Dietzl, W. Maixner, A. Meixner, W. Thomas, J. A. Pospisilik, M. Alenius, M. Kress, S. Subramaniam, P. A. Garrity, H. J. Bellen, C. J. Woolf and J. M. Penninger (2010). “A genome-wide Drosophila screen for heat nociception identifies alpha2delta3 as an evolutionarily conserved pain gene.” Cell 143(4): 628–638.

Newby, M. I. and N. L. Greenbaum (2002). “Sculpting of the spliceosomal branch site recognition motif by a conserved pseudouridine.” Nature Structural Biology 9(12): 958–965.

Ohyama, T., T. Jovanic, G. Denisov, T. C. Dang, D. Hoffmann, R. A. Kerr and M. Zlatic (2013). “High-throughput analysis of stimulus-evoked behaviors in Drosophila larva reveals multiple modality-specific escape strategies.” PLoS One 8(8): e71706.

Ohyama, T., C. M. Schneider-Mizell, R. D. Fetter, J. V. Aleman, R. Franconville, M. Rivera-Alba, B. D. Mensh, K. M. Branson, J. H. Simpson, J. W. Truman, A. Cardona and M. Zlatic (2015). “A multilevel multimodal circuit enhances action selection in Drosophila.” Nature 520(7549): 633–639.

Olesnicky, E. C., D. J. Killian, E. Garcia, M. C. Morton, A. R. Rathjen, I. E. Sola and E. R. Gavis (2014). “Extensive use of RNA-binding proteins in Drosophila sensory neuron dendrite morphogenesis.” G3 (Bethesda) 4(2): 297–306.

Onodera, K., S. Baba, A. Murakami, T. Uemura and T. Usui (2017). “Small conductance Ca(2+)-activated K(+) channels induce the firing pause periods during the activation of Drosophila nociceptive neurons.” Elife 6.

Pan, L., Y. Q. Zhang, E. Woodruff and K. Broadie (2004). “The Drosophila fragile X gene negatively regulates neuronal elaboration and synaptic differentiation.” Curr Biol 14(20): 1863–1870.

Parks, A. L., K. R. Cook, M. Belvin, N. A. Dompe, R. Fawcett, K. Huppert, L. R. Tan, C. G. Winter, K. P. Bogart, J. E. Deal, M. E. Deal-Herr, D. Grant, M. Marcinko, W. Y. Miyazaki, S. Robertson, K. J. Shaw, M. Tabios, V. Vysotskaia, L. Zhao, R. S. Andrade, K. A. Edgar, E. Howie, K. Killpack, B. Milash, A. Norton, D. Thao, K. Whittaker, M. A. Winner, L. Friedman, J. Margolis, M. A. Singer, C. Kopczynski, D. Curtis, T. C. Kaufman, G. D. Plowman, G. Duyk and H. L. Francis-Lang (2004). “Systematic generation of high-resolution deletion coverage of the Drosophila melanogaster genome.” Nat Genet 36(3): 288–292.

Phillips, B., A. N. Billin, C. Cadwell, R. Buchholz, C. Erickson, J. R. Merriam, J. Carbon and S. J. Poole (1998). “The Nop60B gene of Drosophila encodes an essential nucleolar protein that functions in yeast.” Mol Gen Genet 260(1): 20–29.

Ran, F. A., P. D. Hsu, J. Wright, V. Agarwala, D. A. Scott and F. Zhang (2013). “Genome engineering using the CRISPR-Cas9 system.” Nat Protoc 8(11): 2281–2308.

Rangaraju, V., S. Tom Dieck and E. M. Schuman (2017). “Local translation in neuronal compartments: how local is local?” EMBO Rep 18(5): 693–711.

Raychaudhuri, S., L. Niu, J. Conrad, B. G. Lane and J. Ofengand (1999). “Functional effect of deletion and mutation of the Escherichia coli ribosomal RNA and tRNA pseudouridine synthase RluA.” J Biol Chem 274(27): 18880–18886.

Schwartz, S., D. A. Bernstein, M. R. Mumbach, M. Jovanovic, R. H. Herbst, B. X. Leon-Ricardo, J. M. Engreitz, M. Guttman, R. Satija, E. S. Lander, G. Fink and A. Regev (2014). “Transcriptome-wide mapping reveals widespread dynamic-regulated pseudouridylation of ncRNA and mRNA.” Cell 159(1): 148–162.

Tenenbaum, C. M., M. Misra, R. A. Alizzi and E. R. Gavis (2017). “Enclosure of Dendrites by Epidermal Cells Restricts Branching and Permits Coordinated Development of Spatially Overlapping Sensory Neurons.” Cell Rep 20(13): 3043–3056.

Tracey, W. D., Jr. (2017). “Nociception.” Curr Biol 27(4): R129–R133.

Tracey, W. D., R. I. Wilson, G. Laurent and S. Benzer (2003). “painless, a Drosophila gene essential for nociception.” Cell 113(2): 261–273.

Tsubouchi, A., J. C. Caldwell and W. D. Tracey (2012). “Dendritic filopodia, Ripped Pocket, NOMPC, and NMDARs contribute to the sense of touch in Drosophila larvae.” Curr Biol 22(22): 2124–2134.

Turner, H. N., K. Armengol, A. A. Patel, N. J. Himmel, L. Sullivan, S. C. Iyer, S. Bhattacharya, E. P. R. Iyer, C. Landry, M. J. Galko and D. N. Cox (2016). “The TRP Channels Pkd2, NompC, and Trpm Act in Cold-Sensing Neurons to Mediate Unique Aversive Behaviors to Noxious Cold in Drosophila.” Curr Biol 26(23): 3116–3128.

Venken, K. J., J. W. Carlson, K. L. Schulze, H. Pan, Y. He, R. Spokony, K. H. Wan, M. Koriabine, P. J. de Jong, K. P. White, H. J. Bellen and R. A. Hoskins (2009). “Versatile P[acman] BAC libraries for transgenesis studies in Drosophila melanogaster.” Nat Methods 6(6): 431–434.

Venken, K. J. T., K. L. Schulze, N. A. Haelterman, H. Pan, Y. He, M. Evans-Holm, J. W. Carlson, R. W. Levis, A. C. Spradling, R. A. Hoskins and H. J. Bellen (2011). “MiMIC: a highly versatile transposon insertion resource for engineering Drosophila melanogaster genes.” Nature methods 8(9): 737–743.

Vicidomini, R., A. Petrizzo, A. di Giovanni, L. Cassese, A. A. Lombardi, C. Pragliola and M. Furia (2017). “Drosophila dyskerin is required for somatic stem cell homeostasis.” Sci Rep 7(1): 347.

Walcott, K. C. E., S. E. Mauthner, A. Tsubouchi, J. Robertson and W. D. Tracey (2018). “The Drosophila Small Conductance Calcium-Activated Potassium Channel Negatively Regulates Nociception.” Cell Rep 24(12): 3125–3132 e3123.

Wang, C. C., J. C. Lo, C. T. Chien and M. L. Huang (2011). “Spatially controlled expression of the Drosophila pseudouridine synthase RluA-1.” Int J Dev Biol 55(2): 223–227.

Xiang, Y., Q. Yuan, N. Vogt, L. L. Looger, L. Y. Jan and Y. N. Jan (2010). “Light-avoidance-mediating photoreceptors tile the Drosophila larval body wall.” Nature 468(7326): 921–926.

Xu, X., J. L. Brechbiel and E. R. Gavis (2013). “Dynein-dependent transport of nanos RNA in Drosophila sensory neurons requires Rumpelstiltskin and the germ plasm organizer Oskar.” J Neurosci 33(37): 14791–14800.

Ye, B., C. Petritsch, I. E. Clark, E. R. Gavis, L. Y. Jan and Y. N. Jan (2004). “Nanos and Pumilio are essential for dendrite morphogenesis in Drosophila peripheral neurons.” Curr Biol 14(4): 314–321.

Zhong, L., A. Bellemer, H. Yan, H. Ken, R. Jessica, R. Y. Hwang, G. S. Pitt and W. D. Tracey (2012). “Thermosensory and nonthermosensory isoforms of Drosophila melanogaster TRPA1 reveal heat-sensor domains of a thermoTRP Channel.” Cell Rep 1(1): 43–55.

Zhong, L., R. Y. Hwang and W. D. Tracey (2010). “Pickpocket is a DEG/ENaC protein required for mechanical nociception in Drosophila larvae.” Curr Biol 20(5): 429–434.

