## Supplementary figures and images for "Loss of pseudouridine synthases in the RluA family causes hypersensitive nociception in *Drosophila*"

### Figure S1

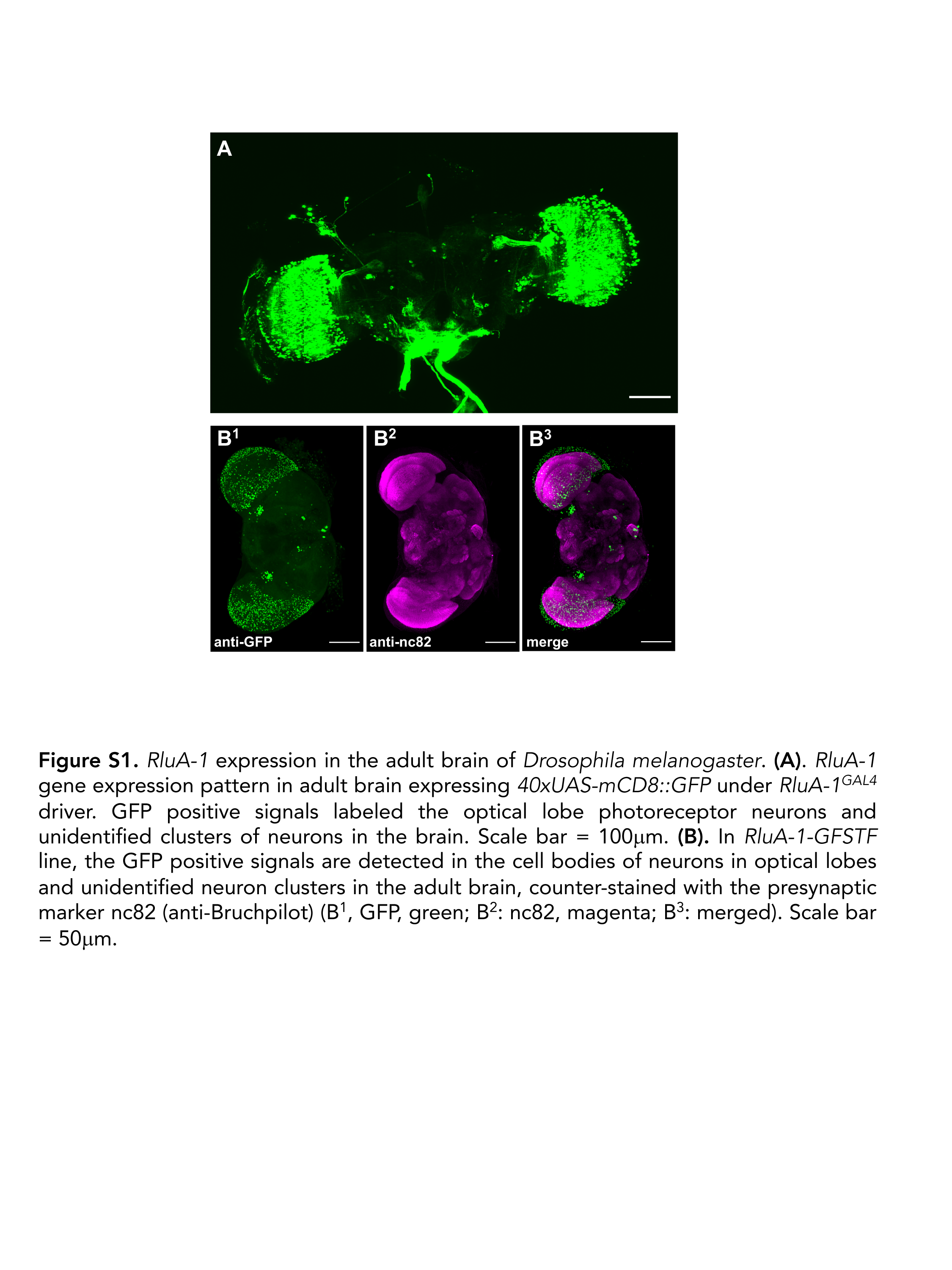

### Figure S3

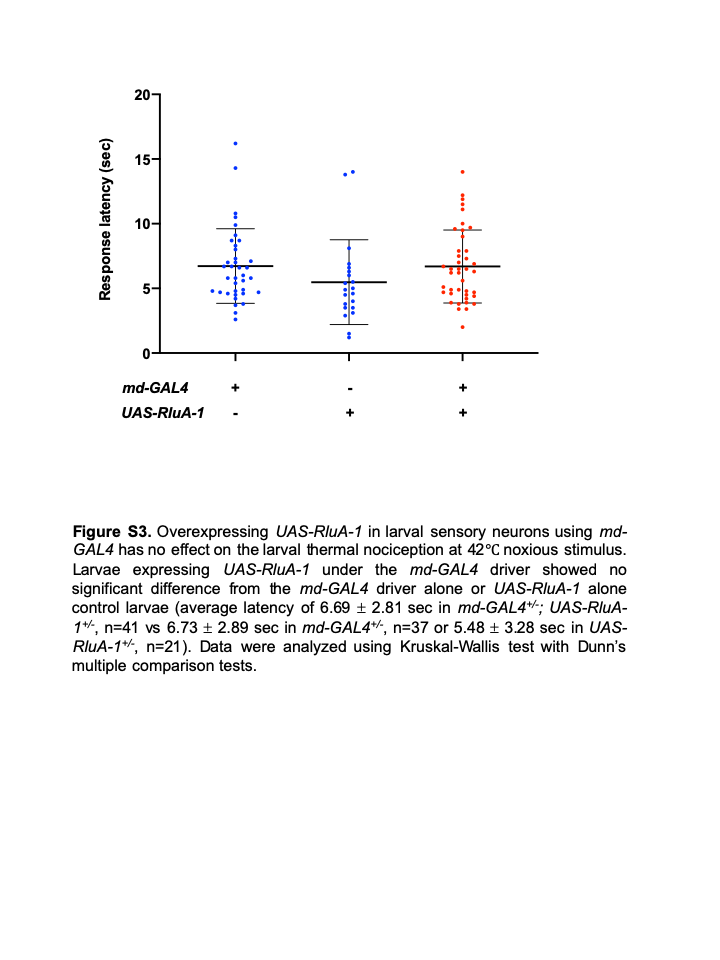

### Figure S5

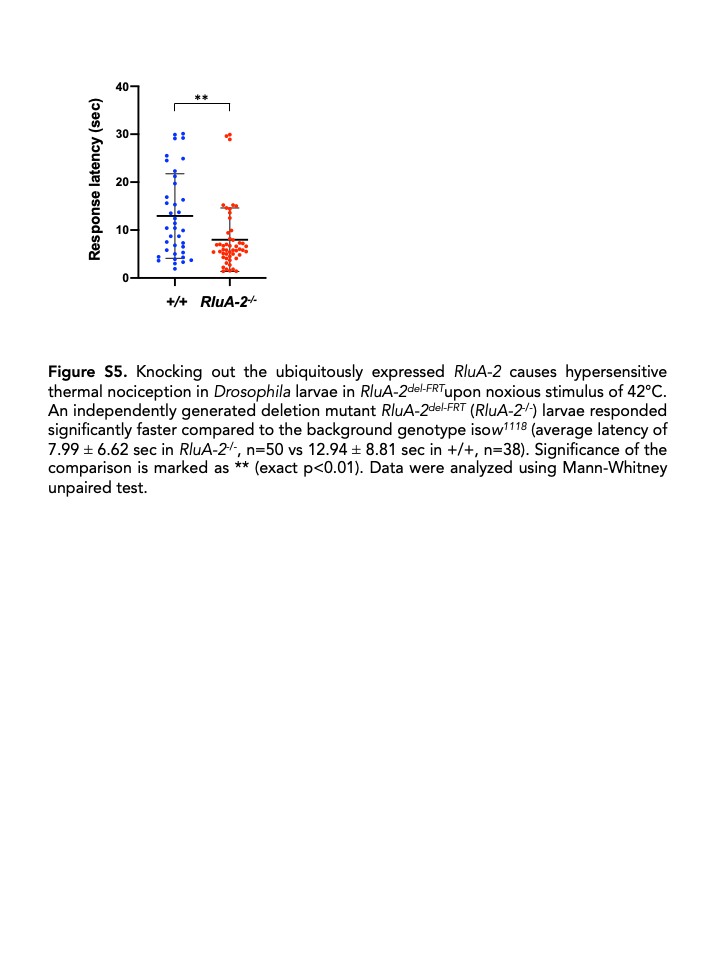
