## Supplementary material for "Loss of pseudouridine synthases in the RluA family causes hypersensitive nociception in *Drosophila*": Figure S2

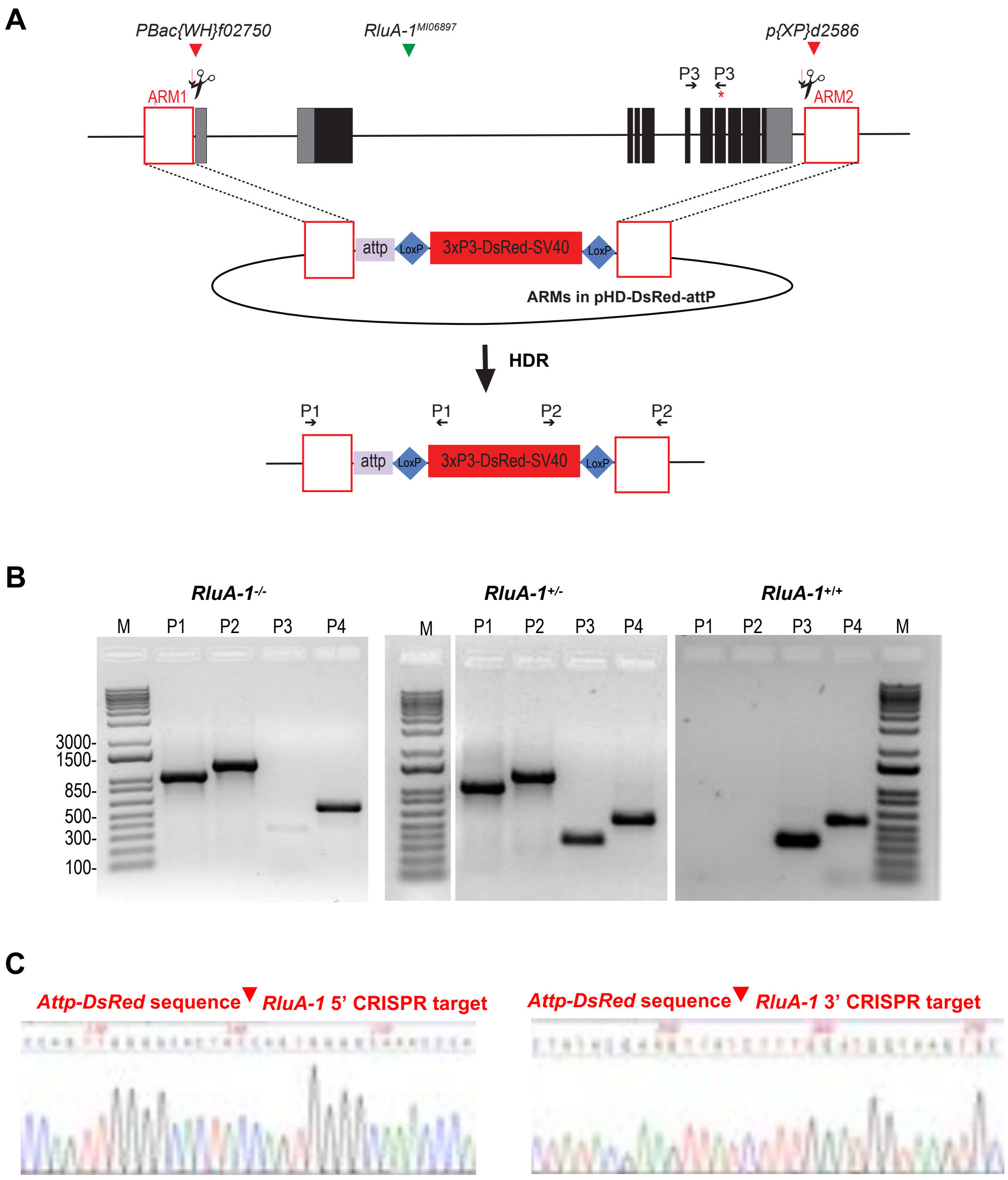

A
PBac{WH}f02750
RluA-1MI06897
p{XP}d2586
P3
P3
ARM1
ARM2
attp
LoxP
3xP3-DsRed-SV40
LoxP
ARMs in pHD-DsRed-attP
HDR
P1
P1
P2
P2
attp
LoxP
3xP3-DsRed-SV40
LoxP
B
RluA-1-/-
RluA-1+/-
RluA-1+/+
M
P1
P2
P3
P4
M
P1
P2
P3
P4
P1
P2
P3
P4
M
3000-
1500-
850-
500-
300-
100-
C
Attp-DsRed sequence
RluA-1 5' CRISPR target
Attp-DsRed sequence
RluA-1 3' CRISPR target

**Figure S2.** Generation and confirmation of the null allele *RluA-1*$^{del\text{-}HDR}$. **(A).** Schematic of *RluA-1* locus and generation of the *RluA-1* deletion allele *RluA-1*$^{del\text{-}HDR}$ with CRISPR/Cas9 induced HDR mediated precise deletion. UTRs are represented as grey boxes, exons as black boxes and introns as black lines, drawn to scale. Homology arms ("ARM1" and "ARM2", indicated as red rectangles) of ~1kb immediately flanking the CRISPR cleavage sites (indicated with the scissor symbols) were cloned into *pHD-DsRed-attP* vector (not drawn to scale). The genomic region between the cleavage sites was replaced with the *attP* ΦC31 docking site and a *3xP3-DsRed* marker flanked by loxP recombination sites. The red triangles represent the locations of the two insertions, *PBac{WH}*$^{f02750}$(+) and *p{XP}*$^{d2586}$(-) used to generate the *RluA-1*$^{del\text{-}FRT}$ allele. The green triangle represents the location of the coding intron localized MiMIC line *RluA-1*$^{MI06897}$ used to generate the *RluA-1*$^{Gal4}$ and *RluA-1-GFSTF* lines. The red asterisk indicates the location of the aspartate residue conserved in all pseudouridine sythases. The approximate locations of primer pairs 1 (P1), 2(P2), and 3(P3) used for PCR confirmation in **Figure S2B** are also shown. A fourth pair of primers (P4) are located on the *RluA-2* locus shown in **Figure S4A** and the presence of the PCR product serves as positive control for integrity of the genomic DNA. **(B).** PCR verification of *RluA-1* deletion allele *RluA-1*$^{del\text{-}HDR}$**. Upper panel**: The resulting deletion mutant allele *RluA-1*$^{del\text{-}HDR}$ over a deficiency *Df(2L)Exel7048* (***RluA-1***$^{-/-}$, left) amplified the expected products from P1 (5' CRISPR target site of *RluA-1* and *attP-DsRed* cassette, P1-F: *RluA-1*-ARM1F, P1-R: *Hsp70*-R, expected product length: 1057bp), P2 (*attP-DsRed* cassette and *RluA-1* 3' CRISPR target site, P2-F: SV40F, P2-R: RluA-1-ARM2R, expected product length: 1311bp), P4 (*RluA-2*, positive control, P4-F:5'-CAACACGGTAGTTTTCATCCTGG-3', P4-R: 5'-CCTCCTCTTCCTTTCAATGGACC-3', expected product length: 562bp) but not from P3 (deleted region containing pseudouridine synthase domain in *RluA-1,* P4-F*: 5'*-GTGACTCCTTGTGACTTACAGGTGC-3', P4-R: 5'-GTTTTCAGAGCACCACACTG-3', expected product length: 341bp). Siblings of *RluA-1*$^{del\text{-}HDR}$ over *CyO* (***RluA-1***$^{+/-}$, middle) amplified all of the expected products from P1, P2, P3 and P4 while the injection line 1 (*vas-Cas9,* ***RluA-1***$^{+/+}$, right) failed to amplify from P1 and P2 but amplified expected products from P3 and P4. "M" represents the 1kb-plus molecular ladder (Invitrogen). **(C)** Sequencing verification of *RluA-1* deletion allele *RluA-1*$^{del\text{-}HDR}$**.** Sanger sequencing chromatographs in *RluA-1*$^{del\text{-}HDR}$ at the 5' CRISPR site (left panel) and 3' CRISPR site (right panel) show the replacement of *RluA-1* genomic sequence with the *attP-DsRed* replacement cassette fragment at the indicated 5' and 3' *RluA-1*-CRISPR target cut sites (marked with red arrowheads).
