## Supplementary material for "Loss of pseudouridine synthases in the RluA family causes hypersensitive nociception in *Drosophila*": Figure S4

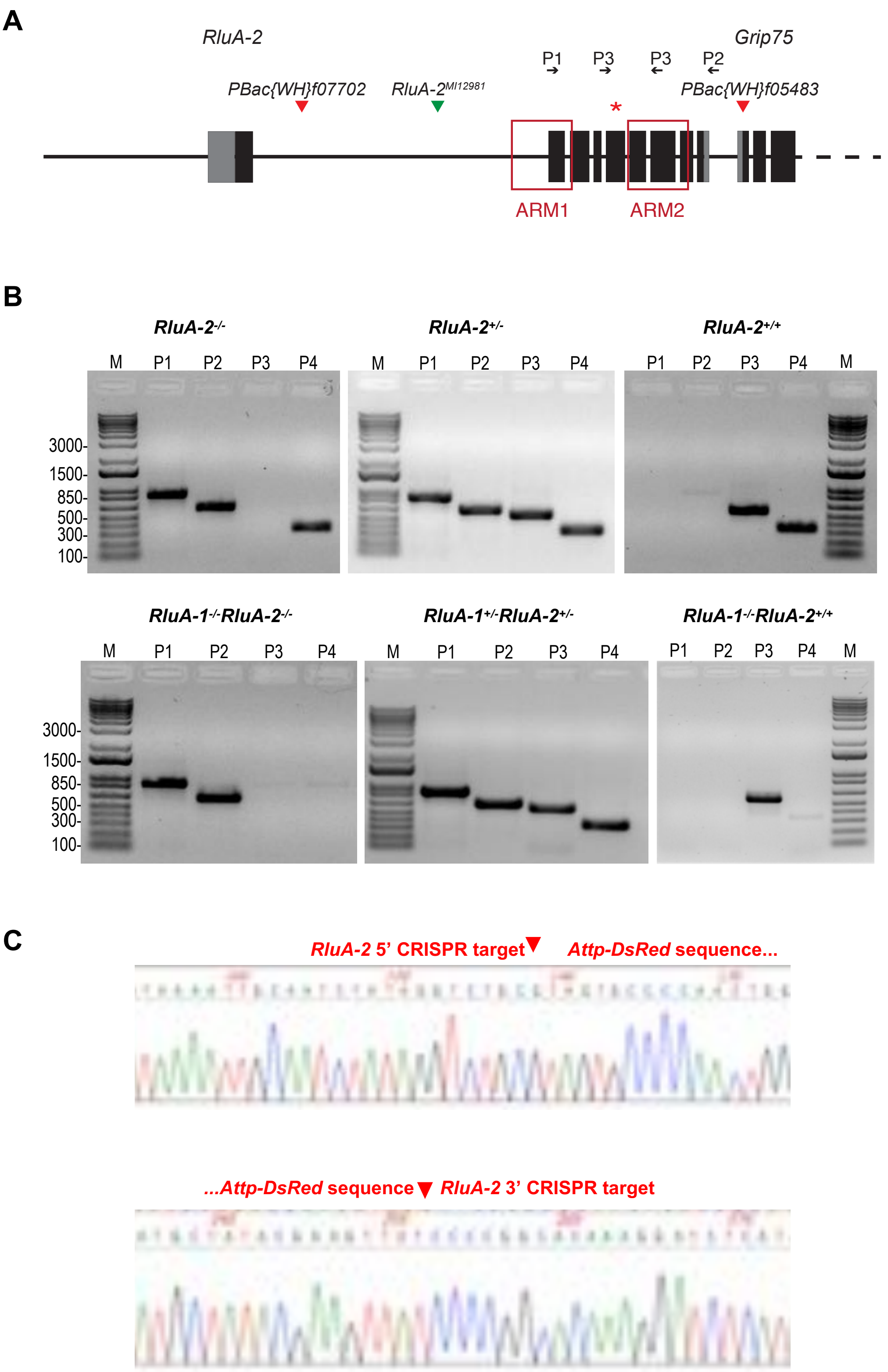
A
RluA-2
Grip75
P1
P3
P3
P2
PBac{WH}f07702
RluA-2^MI12981
PBac{WH}f05483
*
ARM1
ARM2
B
RluA-2-/-
RluA-2+/-
RluA-2+/+
M
P1
P2
P3
P4
3000-
1500-
850-
500-
300-
100-
RluA-1-/-RluA-2-/-
RluA-1+/-RluA-2+/-
RluA-1-/-RluA-2+/+
C
RluA-2 5' CRISPR target
Attp-DsRed sequence...
...Attp-DsRed sequence
RluA-2 3' CRISPR target

**Figure S4.** Generation and confirmation of *RluA-2* single deletion allele *RluA-2$^{del\text{-}HDR}$* and double deletion allele *RluA-1$^{del\text{-}HDR}$ RluA-2$^{del\text{-}HDR}$*. (**A**). Schematic of *RluA-2* gene locus. UTRs are represented as grey boxes, exons as black boxes and introns as black lines (drawn to scale). Partial sequence of the downstream gene Grip75 was also shown. Homology arms ("ARM1" and "ARM2") immediately flanking the CRISPR cleavage sites are marked as red rectangles. The red triangles mark the locations of the two insertions, *PBac{WH}f07702* and *PBac{WH}f05483* used to generate the *RluA-2$^{del\text{-}FRT}$* allele. The green triangle marks the location of the coding intron localized MiMIC line (*RluA-2*$^{MI12981}$) used for generating the *RluA-2-GFSTF* line. Red star (*) indicates the location of the aspartate residue conserved in all pseudouridine sythases. The approximate locations of primer P1-F, P2-R, and P3 used for PCR confirmation (**Figure S4B**) were also shown. P1-R and P2-F are located on *attP-DsRed* cassette as shown in **Figure S2A** while the P3 primers located on *RluA-1* as shown in **Figure S2A** are used as P4 in **Figure S4B**. (**B**). PCR verification of single mutant *RluA-2$^{del\text{-}HDR}$* and double mutant *RluA-1$^{del\text{-}HDR}$ RluA-2$^{del\text{-}HDR}$*. **Upper panel**: Single mutant allele *RluA-2$^{del\text{-}HDR}$* over the deficiency *Df(2L)Exel7048 ("RluA-2$^{-/-}$")* amplified the expected products from P1 (5' border of *RluA-2* and *attP-DsRed* cassette, P1-F: 5'-TGTCTATGTTTGCTTCCTGGCT-3', P1-R: Hsp70-R, expected product length: 866bp), P2 (3' border of *attP-DsRed* cassette and *RluA-2*, P2-F: SV40-F, P1-R:5'-CACGACGATAAATAATACTCCACG-3', expected product length: 744bp), and P4 (*RluA-1* pseudouridine-synthase domain containing region, 341bp), but not from the deleted region in *RluA-2* with P3 (562bp). Siblings of *RluA-2$^{del\text{-}HDR}$* over CyO (*"RluA-2$^{+/-}$"*) amplified all of the expected products from P1, P2, P3 and P4 while the injection line 2 (*Nos-Cas9, "RluA-2$^{+/+}$"*) failed to amplify from P1 and P2 but amplified the expected products with P3 and P4. **Bottom panel**: Double mutant alleles *RluA-1$^{del\text{-}HDR}$RluA-2$^{del\text{-}HDR}$* over the deficiency *Df(2L)Exel7048 ("RluA-1$^{-/-}$RluA-2$^{-/-}$")* amplified the expected products from P1 (5' border of *RluA-2* and *attP-DsRed* cassette, 866bp), P2 (3' border of *attP-DsRed* cassette and *RluA-2*, 744bp), but not from the deleted region in *RluA-2* with P3 and deleted region in *RluA-1* with P4. Siblings of *RluA-1$^{del\text{-}HDR}$RluA-2$^{del\text{-}HDR}$* over CyO (*"RluA-1$^{+/-}$RluA-2$^{+/-}$"*) amplified all of the expected products from P1, P2, P3 and P4 while the injection line 3 (*RluA-1*; *Nos-Cas9, "RluA-1$^{-/-}$RluA-2$^{+/+}$"*) failed to amplify from P1 and P2 and P4 but amplified the expected products with P3. M represents the 1kb-plus molecular ladder (Invitrogen). **(C)**: Sequencing verification of single mutant *RluA-2$^{del\text{-}HDR}$* and double mutant *RluA-1$^{del\text{-}HDR}$ RluA-2$^{del\text{-}HDR}$*. Sanger sequencing chromatographs at the 5' CRISPR site (upper) and 3' CRISPR site (bottom) show *RluA-2* genomic sequences are replaced with the *attP-DsRed* replacement cassette fragment in single mutant *RluA-2$^{del\text{-}HDR}$* or the double mutant *RluA-1$^{del\text{-}HDR}$ RluA-2$^{del\text{-}HDR}$* at the indicated 5' and 3' *RluA-1*-CRISPR target cut sites (marked with red arrow heads).
